# Unveiling the Conformational Dynamics of the Histone Tails Using Markov State Modeling

**DOI:** 10.1101/2025.01.16.633411

**Authors:** Rutika Patel, Sharon M. Loverde

**Author notes:** Sharon M. Loverde **Email:**. PNAS strongly encourages authors to supply an ORCID identifier for each author. Do not include ORCIDs in the manuscript file; individual authors must link their ORCID account to their PNAS account at www.pnascentral.org. For proper authentication, authors must provide their ORCID at submission and are not permitted to add ORCIDs on proofs. **Author Contributions:** R.P parameterized and conducted molecular dynamics simulations, analyzed data, prepared all the figures, and wrote the paper. The study and paper writing are supported by S.M.L. S.M.L also supervised the project. **Competing Interest Statement:** None. **Classification:** Biophysics and Computational Biology, Physical Science.

## Abstract

Biomolecules predominantly exert their function through altering conformational dynamics. The nucleosome core particle (NCP) is the fundamental unit of chromatin. DNA with ∼146 base pairs wrap around the histone octamer to form a nucleosome. The histone octamer is comprised of two copies of each histone protein (H3, H4, H2A, and H2B) with a globular core and disordered N-terminal tails. Epigenetic modifications of the histone N-terminal tails play a critical role in the regulation of chromatin structure and biological processes such as transcription and DNA repair. Here, we report all-atomistic molecular dynamics (MD) simulations of the nucleosome at microsecond timescales to construct Markov state models (MSMs) to elucidate distinct conformations of the histone tails. We employ the time-lagged independent component analysis (tICA) to capture their essential slow dynamics, with k-means clustering used to discretize the conformational space. MSMs unveil distinct states and transition probabilities to characterize the dynamics and kinetics of the tails. Next, we focus on the H2B tail, one of the least studied tails. We show that acetylation increases secondary structure formation, with an increase in transition rates. These findings will aid in understanding the functional implications of tail conformations in nucleosome stability and gene regulation.

**Significance Statement:** The nucleosome is the fundamental repeat unit of chromatin, composed of a histone octamer and DNA. Each histone has a globular core and N-terminal tail regions. N-terminal tails are major sites for post-translational modifications, chromatin structure, and gene regulation. Using all-atom molecular dynamics simulations of the nucleosome to examine the dynamics of histone tails and analyze the effects of one of the histone modifications, such as acetylation on histone tails. We explore tail dynamics and kinetics using Markov State Models (MSMs) to gain insight into histone tail structural changes. Acetylation of the H2B tail shows an increased dynamics of the tail based on transition rates. Our study characterizes transition rates of all histone tails that can influence nucleosome plasticity and gene regulation.

## Introduction

In eukaryotic cells, DNA is packaged inside the nucleus through a hierarchical structure involving the nucleosome core particle (NCP). Nucleosomes are the fundamental repeat units of chromatin(1). Each NCP comprises a histone octamer around which approximately 147 base pairs of DNA are wrapped. Two copies of each of the histones H3, H4, H2A, and H2B assemble to make the histone octamer. Together with histone H1 and linker DNA, they further assemble into a higher-order compact and dynamic chromatin structure(1-5). Histones have a globular core region and flexible N-terminal tails that protrude from the histone core. The NCP is stabilized by electrostatic interactions between the negatively charged DNA-phosphate backbone and positively charged histone residues, such as lysine (Lys, K) and arginine (Arg, R) (6, 7). Histone tails are major sites of post-translational modifications (PTMs) such as acetylation, methylation, and phosphorylation, which influence chromatin structure and gene regulation for various biological processes, including DNA repair, replication, and gene expression. PTMs can alter the highly ordered chromatin structure to allow protein modulators to access the DNA (8, 9). Histone tails are intrinsically disordered in nature. However, the tails are dynamic and can still transiently form secondary structures (10, 11).

The histone N-terminal tails perform several critical biological functions, including inter-nucleosome contacts (9), nucleosome stability and dynamics, DNA accessibility, nucleosome sliding, and coordinating various epigenetic pathways in time. The tails also speed up the search for nucleosome targets to ease their interactions and are involved in DNA unwrapping (12). The histone tails are highly positively charged and interact with the negatively charged DNA-phosphate backbone by forming salt bridges. Any changes in the tails via PTMs can perturb these interactions (12, 13). Each histone tail that protrudes from the histone core has a unique sequence; the tails are further distinguished by their positioning with respect to the DNA superhelical locations (SHLs). The H3 N-terminal tails (1-43 residues) are located near the DNA entry/exit regions, and they extend between the DNA gyres near the SHL±7 regions. The H3 tails are usually seen to collapse onto the DNA rather than extend. Compared to the H3 tails that extend near the entry/exit regions of the DNA, the H4 N-terminal tails (1-23 residues) are located on the face of NCP, extending from the core near the SHL±2 regions. Like the H3 tails, the H4 tails mostly collapse onto the DNA. The H2A N-terminal tails (1-15 residues) are positioned around the face of the nucleosome like the H4 tails, but they are further away from the dyad and located near the SHL±4 regions. The H2B N-terminal (1-30 residues) tails protrude between the two DNA gyres, and they come into contact with the DNA around the SHL±5 and SHL±3 regions. The H2B tails are also located very close to the H2A N-terminal tails and away from the dyad. The histone tails cover nearly the entire DNA and make contact with nearly every SHL region. The tails also play an essential role in DNA breathing and unwrapping (14).

Small-angle-X-ray scattering (SAXS) and fluorescence resonance energy transfer (FRET) studies have demonstrated that removing the H3 tail destabilizes the NCP, leading to unwrapping. Also, the removal of the H4 tails causes DNA to be tightly bound to the histone octamer, indicating that tails distinctly influence unwrapping (15). Acetylation increases nucleosome unwrapping, which has been shown by a single-molecule FRET study (16). Several circular dichroism (CD) and Nuclear Magnetic resonance (NMR) studies have characterized the secondary structure conformations of the histone tails (17). An NMR study by Kim *et al (18)* has shown that acetylation of H3, H4, H2A, and H2B histone tails causes subtle changes in NCP dynamics. The tail dynamics are observed at picoseconds to nanoseconds timescales. There is an increase in the motions of the acetylated tails and DNA accessibility for regulatory proteins (18). In addition, other NMR studies of the NCP have provided insights into tail interaction with proteins (19-22). NMR studies have demonstrated that the H3 and H4 tails decrease the compaction of nucleosomal arrays, as acetyl tail dynamics promotes the binding of regulatory proteins (20). Acetylation of one of the H4 tail residues can enhance the acetylation rate for the H3 tail, as suggested by NMR studies (21). All the above MD and biophysical methods studies have characterized the histone tail dynamics and effects of acetylation, but a more detailed characterization of the transient conformational states of the tails, and the rates between these states has yet to be performed.

Molecular dynamics (MD) simulations have elucidated some aspects of nucleosome dynamics; however, the kinetics of these processes can be further outlined. Earlier studies of all-atom MD simulations have characterized DNA breathing (23-25), effects of PTMs of histone H3 and H4 tails that decrease tail-DNA contacts (26-32), and the conformational rearrangement of the N-terminal histone tails. MD simulations have also characterized nucleosome stability based on the H3 and the H2B tail—DNA interactions (33-36), including the role of the histone H2B and H2A tails, and DNA loop formation. Previous studies conducted by our laboratory included DNA partial unwrapping of the 1KX5 NCP system and wave-like motion promoted by stabilizing effects of H2A and destabilizing effects of H2B tails at microsecond timescales (37) (23). Another study from our lab demonstrated the motions of the Widom-601 and alpha satellite palindromic (ASP) NCP DNA sequences at longer microsecond timescales that show different pathways for the two sequences, with one showing loop formation and another showing large-scale breathing. The motion and contact of H2A and H2B tails play a critical role in loop formation. The N-terminal H3 tail plays a key role in the breathing motion of the DNA (38). Thus, all-atom MD simulations are useful for characterizing NCP structure and dynamics.

One of the well-characterized PTMs is lysine acetylation of the histone N-terminal tail residues. Lysine acetylation neutralizes the positive charge of lysine by replacing it with acetyl (-CO-CH_3_) group. Histone acetyltransferases (HATs) transfer the acetyl group from acetyl-Coenzyme A (acetyl-CoA) to the ε-amino group of side chain of lysine residue that neutralizes the positive charge of lysine residues (18). Epigenetic modifications are associated with various diseases, including cancer, neurological disorders, and inflammatory diseases. One of the histone proteins, H2B, is associated with transcription activation (39-41), DNA repair (42), and cancer. In addition, one of the tumor repressor proteins, p14ARF, is associated with H2B acetylation at Lys 5, 12, 15, and 20 tail residues. Our previous molecular dynamics (MD) simulation studies showed that the H2B N-terminal tails promotes the outward stretching of SHL-5 region of DNA in NCP complex in DNA unwrapping (37). Our recent study on H2B tail acetylation also revealed that acetylation makes the tail more dynamic and increases helices as well as decreases DNA-H2B tail contacts. It also showed that acetylation rearranges the secondary structure of the H2B tail, making it more helical and changing the conformational space of the tail based on the principal component analysis (PCA) (34). Histone H2B tail dynamics upon acetylation is a critical factor in biological processes, yet its kinetics remains to be elucidated.

Here, we use Markov State Modeling (MSM) to gain an understanding of the dynamics and conformational space of histone tails in the NCP system based on molecular dynamics (MD) simulation. Conformational transitions of the histone tails are vital to gene regulation. MSM of molecular kinetics, which approximates the long-term dynamics of molecular systems over a discretized conformational space, has gained extensive application in recent years (43, 44). MSMs are used for kinetic analysis by modeling a molecular system as a memoryless transition network, where the probability of transitioning to a future state depends only on the present state, not on the system’s past states. A MSM can describe the entire dynamics of the system. A MSM is an *n x n* square matrix known as a transition probability matrix where the configuration space spanned by the system is divided into *n* states. The transition probability matrix is characterized by *n* states and by the lag time τ at which the state of the system was recorded. The transition state populations and conditional pairwise transition probabilities can be obtained from this matrix. The resulting models are called transition MSM networks. The transition probabilities yield kinetic information as well as possible transition state pathways (TPT) between the states (44).

The dynamic transition between metastable conformational states, such as opening and closing of the SARS-Cov-2 spike protein complex, translocation of RNA polymerases on the DNA template during transcription, etc, are essential to exert their biological functions (45, 46). There are several MSM and related techniques that have been used to study protein folding (47-51) and dynamics (52), protein conformational changes (53-56) and protein-ligand binding (57-59). Histone tails undergo secondary structure conformation changes that include the tail folding into and extended into random coil structures. A previous H3 tail MSM study considered only H3 tail without the DNA of the NCP complex and showed that the H3 tail undergoes conformational changes at nanoseconds timescales (60).

In this study, we perform all-atomistic molecular dynamics (MD) simulations of the NCP at physiological 0.15 M salt concentrations. We run two sets of simulations with one 0.15 M unacetylated (WT) and 0.15 M acetylated H2B N-terminal tails (ACK) systems. Only the H2B tail’s residues K5, K12, K15, and K20 residues are acetylated to study the effects of acetylation on the kinetics of the histone tails. The H2B tail is considered for this as we mentioned that H2B tail in critical yet not well studied, and our previous H2B tail acetylated study shows changes in the conformational space upon acetylation of H2B tails. We build a MSM of WT H3, H4, H2A, H2B, and the ACK H2B N-terminal tails. We perform feature selection using the VAMP-2 scoring method and selected features as a combination of backbone torsions and pairwise distances. Further, we perform dimensionality reduction using TICA for all histone tails that reveal distinct minima. The kinetic information of the tail is extracted based on MFPT between different transition states of the tails in nanoseconds. In addition, the MFPT between the WT H3, H4, H2A, and H2B tails show that the H2A tail, being the shortest among other tails, has the shortest MFPT compared to the other tails. The MFPT of the acetylated H2B tails is slightly faster than the WT H2B tails. Overall, our results establish a clear understanding of histone tails kinetics that add insight into nucleosome dynamics and other biological functions.

## Results

### Nucleosome Structure and MSM Overview

Here, we perform multiple molecular dynamics (MD) simulations of the nucleosome core particle (NCP) of the 1KX5(37) system. We perform simulations of the wild-type (WT) and H2B tail lysine-acetylated (ACK) systems with four lysine K5, K12, K15 and K20 acetylated at 0.15 M salt concentration each for 6 µs. We analyze the kinetics of WT H3, H4, H2A, and H2B tails as well as acetylated (ACK) H2B N-terminal tails by constructing the Markov state models (MSMs). The structure of the NCP (PDB ID: 1KX5(37)) system consists of DNA and histone proteins, as shown in **Fig. 1A**. The NCP consists of 147 DNA base pairs wrapped around two copies of the histone proteins H3, H4, H2A, and H2B. The crystal structure of the 1KX5 system was characterized by Davey *et al.* (37) at resolution of 1.9 Å. This system has the N-terminal tails resolved in the crystal structure for the histone octamer. The DNA of the NCP has superhelical location (SHL) regions consisting of approximately ten base pairs **(Fig. 1A)**. The orientation of the DNA base pairs of the NCP is represented relative to the central base pair, known as SHL zero. The SHL is given where the major groove faces the histone octamer. The first SHL is SHL 0 at the NCP dyad, and the last is SHL±7.

**Fig. 1.**
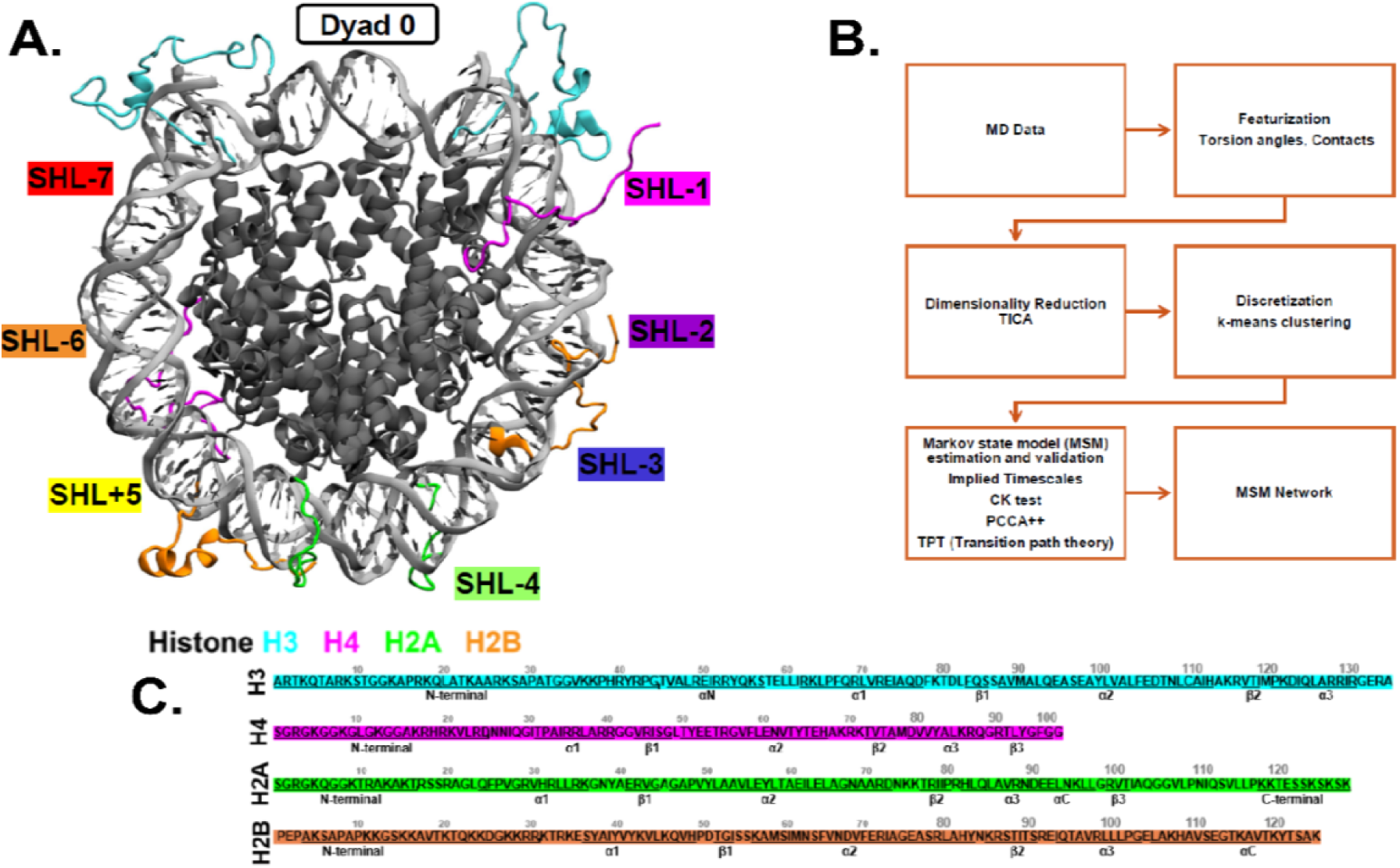
Overview of Nucleosome Core Particle (NCP) Structure and MSM Workflow. **(A)** The crystal structure of NCP (PDB ID: 1KX5) consists of 147 DNA base pairs wrapped around two copies of the histone proteins H3 (cyan), H4 (magenta), H2A (green), and H2B (orange). The super helix locations (SHLs) of DNA are indicated with different colors on the outer side of NCP. **(B)** MD simulations are performed, and MD trajectories are processed to select relevant features as an input. Dimensionality reduction is performed using the selected features. The dimensionality reduction data is discretized to obtain microstates using k-means clustering algorithm. Microstates are assigned to macrostates using PCCA algorithm. MSM estimation is done based on implied timescales (Its) and validation through CK test. MSM network plot can be generated as a graph where the nodes present each MSM states. **(C)** The sequence of histone proteins H3, H4, H2A, and H2B with N-terminal residues are shown for all histone proteins. Out of all histones, only histone H2B N-terminal tail’s four lysine residues are acetylated.

Each histone protein of the NCP has N-terminal tail regions, which are more prone to post-translational modifications. The N-terminal tails do not have a specific tertiary structure, but they undergo changes in secondary structure conformation. Here, we acetylated four lysine residues (K5, K12, K15, K20) of both H2B N-terminal tails by adding the acetyl group, neutralizing the positive charge of these lysine residues. Other H3, H4, and H2A N-terminal tails are not acetylated. Our previous study, Patel *et* al (34)., has shown that the histone tails undergo conformational changes by transitioning from one state to another throughout MD simulations, and the acetylation of H2B tail changes the conformational space of the tail. Principal component analysis (PCA) of the H2B N-terminal tails for WT and ACK systems have a free energy landscape with distinct basins belonging to specific N-terminal tail conformations. In another study from our lab, Khatua *et al (38)* has also shown the PCA of the H3 N-terminal tails for WT systems with distinct conformations. As the N-terminal tails undergo distinct transient conformational changes, the dynamic properties of the histone tails, such as the kinetics of transitions among different conformations, can provide insight into these conformational changes. Therefore, in this study, we use Markov State Models (MSM) to gain further understanding of the dynamics of the histone tails in the WT and the H2B N-terminal tail acetylated systems. H2B acetylation is chosen as our previous study based on H2B acetylation showed an increase in acetylated tail dynamics and underwent secondary structural changes compared to the WT at microsecond timescales. In addition, H2B is a critical gene regulator in cancer and nucleosome dynamics, yet it is not well characterized. Here, we show the kinetics of WT H3, H4, H2A, H2B as well as acetylated H2B tails, to see the effects of acetylation on kinetics of the H2B N-terminal tails.

The workflow for the MSM of the MD simulations consists of the steps shown in **Fig. 1B**. First, the features from the raw MD data can be extracted that represent the dynamics of the system. Examples of the features include torsion angles, contacts, pairwise distance, coordinates of the protein backbone, etc. Then, the high dimensional feature space is transformed into a reduced dimensional space using methods such as principal component analysis (PCA), time-lagged independent component analysis (tICA) to identify key feature components. For MSM analysis, one of the dimension reduction techniques is tICA, which constructs linear combinations of the features to identify the reaction coordinates of the slow timescale processes. Further, the MD trajectories are discretized with the tICA space into a state decomposition of the system using the *k-*means algorithm. The k*-*means clustering algorithm segments the reduced tICA space into a number of cluster centers that group similar conformations in the same cluster centers. These clusters are known as microstates of the system. Next, MSM estimation is performed to choose a lag time (τ) long enough to ensure Markovian dynamics. Therefore, implied timescales (ITS) as a function of τ can be plotted to select the MSM lag time. The microstates are coarse-grained into a smaller number of macrostates using the PCCA+ algorithm, and then the MSM validation is performed with the CK test. The CK test can show that the MSM of our system agrees well with the MSM estimated with longer lag times. The transitions between the states are counted and converted to transition probabilities. The transition probabilities include all transitions between every state, including self-transitions. These probabilities can be visualized on the MSM network plot. Other kinetic analysis, such as transition path theory (TPT), can be also performed to observe the probability flux from initial state to final state. In addition, the rates can be measured by calculating the mean first passage time (MFPT) of the MSM.

### Feature selection and dimensionality reduction of Histone N-terminal tails

In this work, various analyses are performed based on the explicit solvent simulations of the entire nucleosome core particle (NCP), which is composed of DNA with a histone core and N-terminal tails. To initiate the Markov state modeling, the first step is to extract the features from MD trajectories that best represent the conformational states of the histone N-terminal tails. To approximate slow dynamics, the set of input features from each instantaneous configuration of the system must be defined. As we observed earlier in our previous study (34) and during our simulations, histone N-terminal tails undergo secondary structure changes. Therefore, we have selected the internal coordinates of histone tails as backbone torsions, as the tails undergo several secondary structure changes, and pairwise distance between C_α_ carbon of amino acid residues of the tails, as combined input features to build the MSM. In addition, we have calculated a scalar score obtained by using the variational approach to Markov processes (VAMP) to compare certain features that capture the slow dynamics modes of histone tails. We calculate the VAMP-2 score that maximizes the kinetic variance present in the selected features. We compute the VAMP-2 score for different features for different lag times and find relative rankings of the different features as a function of lag time **(*SI Appendix*, Fig. S1).** We find that the combination of backbone torsions and pairwise distance contains more kinetic variance than other features such as torsions, distance, and inverse distance. This suggests that the combination of backbone torsions and pairwise distance are the best features evaluated to build the MSM. A previous study about protein folding also constructs MSM by using similar features — combined backbone torsions and pairwise distance as input features (61). As there are two copies of histone tails in the nucleosome, we have analyzed both copies for H3, H4, H2A, H2B and ACK H2B histone tails to construct the MSM.

Next, we perform tICA (Time-lagged -Independent Component Analysis) to reduce the dimension from the feature space, which typically contains many degrees of freedom, to a reduced dimensional space that can be discretized. tICA is computed on the features to yield reduced dimensional space for all histone tails in the WT system and H2B acetylated tails using PyEMMA (62). The free energy projected on the leading two independent components (ICs) exhibits several distinct minima for the histone tails **(Fig. 2 *A-E*, *SI Appendix*, Fig. S2 *A-E*)**. As there are two copies of histone tails for each histone protein, we have provided data for all the histones Tail-1 in the main **(Fig. 2 *A-E*)** and Tail-2 in *SI appendix **(SI Appendix*****, Fig. S2 *A-E*)**. Based on several minima, we assume that our tICA transformed features describe more than one metastable process in histone tails. After tICA provides dimensionally reduced data of the MD simulation, these reduced data can facilitate the decomposition of the histone tails system into the discrete Markovian states required for MSM estimation. We use the *k*-means algorithm to segment the tICA space into approximately *k=*200 cluster centers for almost all histone tails, except one of the histone tails, which required 75 cluster centers. The *k-*means algorithm segments the reduced tICA space into the number of cluster centers that group similar conformations of the WT H3, H4, H2A, H2B and the ACK H2B histone tails. These clusters are known as microstates.

**Fig. 2.**
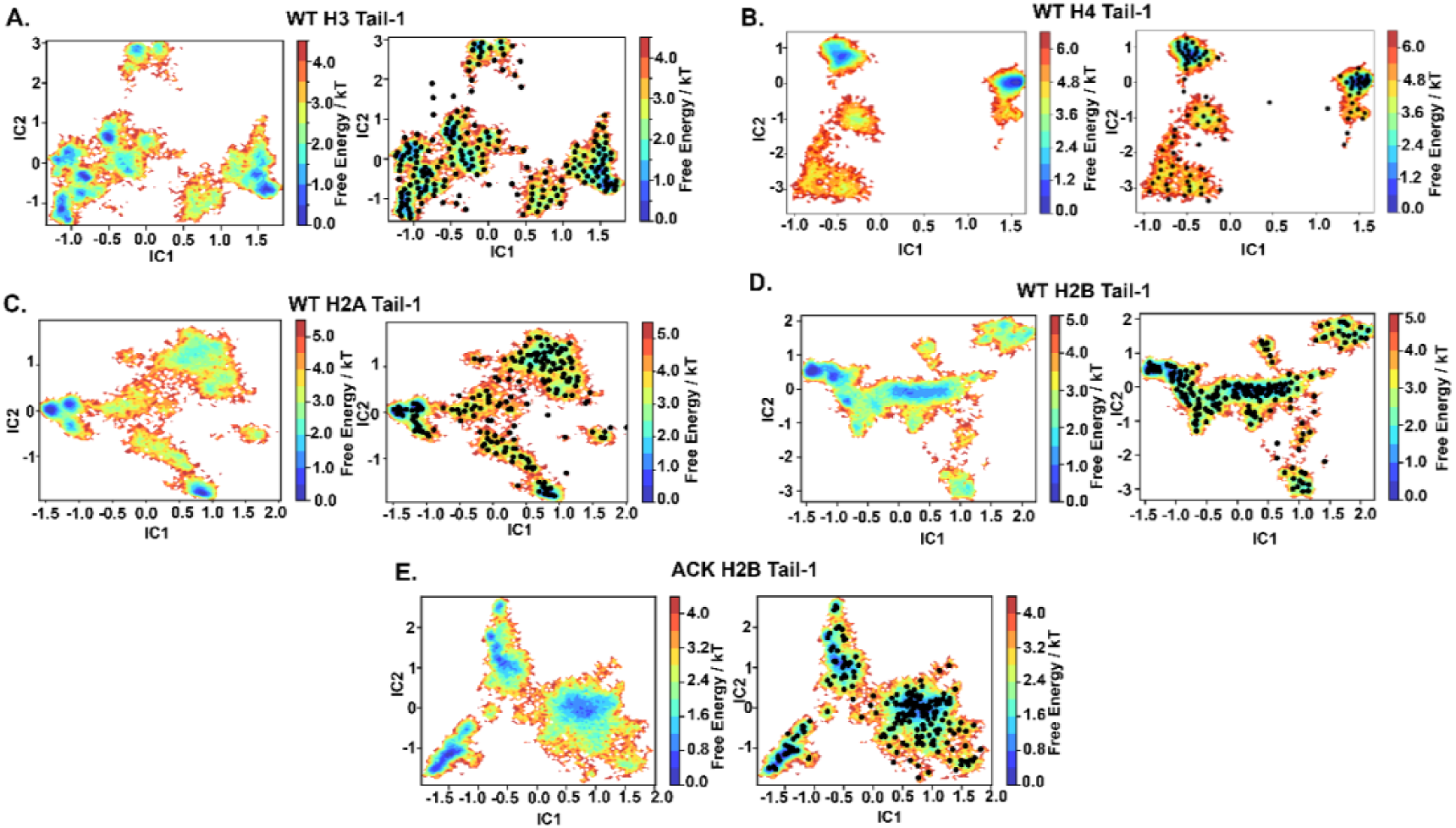
Dimensionality Reduction using Time Independent Component Analysis (TICA). The free energy surface and *k-*means clustering visualizations are projected onto the leading two Independent Components (ICs) for WT **(A)** H3 tail-1 **(B)** H4 tail-1 **(C)** H2A tail-1 **(D)** H2B tail-1 and **(E)** ACK H2B tail-1. The *k-*means clustering provides microstates for each tails based on their conformational space.

### Markov State Model estimation and validation of Histone N-terminal tails

To estimate the MSMs based on our reduced tICA space, the criterion is that the implied timescales (ITs) are approximately constant as a function of lag time τ. Subsequently, the smallest possible lag time τ can be chosen that satisfies this criterion. In the NCP system, the histone N-terminal tails H3, H4, H2A, H2B, and ACK H2B for Tail-1 show five slow processes, and these processes are constant for lag times above 5 ns **(Fig. 3A-E)**. Therefore, we can now estimate a MSM with lag time τ at 5 ns and can perform a validation test. Similarly, for Tail-2 ***(SI Appendix*, Fig. S3 A-E)**, for all histone N-terminal tails H3, H4, H2A, H2B and ACK H2B show five slow processes that are constant at different lag times above 5 ns.

**Fig. 3.**
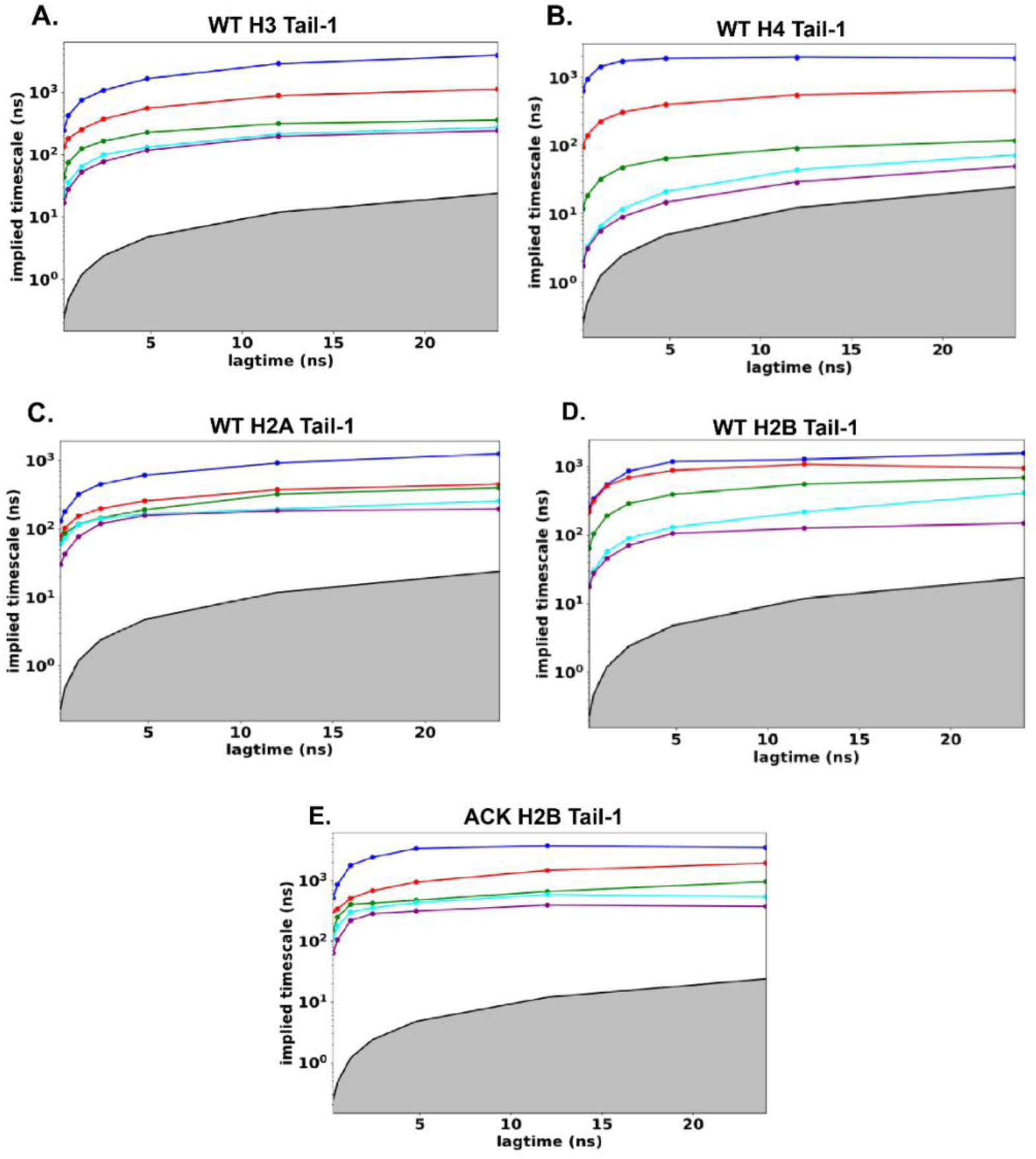
Implied Timescales of Histone N-terminal tails. The implied timescales (ITs) are associated with the five slowest processes for WT **(A)** H3 tail-1 **(B)** H4 tail-1 **(C)** H2A tail-1 **(D)** H2B tail-1 and **(E)** ACK H2B tail-1. The implied timescales plots show Markov processes with different lag times. The solid line corresponds to the maximum likelihood. The blackline with grey shaded area indicates the timescale horizon below which the MSM cannot resolve processes.

We perform a Chapman-Kolmogorov (CK) test to validate the MSM for the histone tails. Before performing the CK test, an appropriate number of metastable states are chosen for the histone tails **(*SI Appendix* Fig. S4-S8)**. For the CK test of the histone tails, the predictions from our MSM (blue-dash line in Figure S4-S8) agree well with the MSM estimated (solid line in Figure S4-S8). The CK tests are performed at the chosen lag time based on the implied timescales that can predict the long-timescale behavior for all histone tails. The CK test for WT H3 **(*SI Appendix* Fig. S4)** shows four metastable states for Tail-1 and Tail-2, and the prediction from our MSM agrees well with the estimated MSM. Similarly, for histone tail H4, both Tail-1 and Tail-2 CK test shows good agreement for four and five metastable states, respectively **(*SI Appendix* Fig. S5)**. For both H2A tails, four metastable states agree with well with our predicted and estimated MSM **(*SI Appendix* Fig. S6)**. For WT H2B Tail-1 and 2, four metastable states are in good agreement as per the CK test, whereas, for acetylated H2B tails, five metastable states for Tail-1 and 2 are in good agreement according to the CK test **(*SI Appendix* Fig. S7-S8)**. As all tails show excellent agreement with our predicted MSM and estimated MSM for the CK test, this confirms the choice of metastable states and the lag time.

As mentioned earlier, *k*-means clustering is used to obtain microstates for the histone N-terminal tails; these microstates of histone tails can be clustered to coarse-grain 200 microstates into appropriate four to five macrostates depending upon the histone tails conformational space. The probability of each microstate belonging to metastable macrostates provides the “membership” of each microstate into a few macrostates. For the WT H3 tails and H2A tails, PCCA+ has identified four macrostates. For the H4 Tail-1 and Tail-2, PCCA+ has identified four and five macrostates. For the WT H2B tails, PCCA+ has identified four macrostates; whereas, for the acetylated H2B tails, PCCA+ has identified five macrostates **(*SI Appendix* Figure S9-S10)**. Next, the MSM is constructed between these macrostates for each of the histone tails.

### MSM Analysis and Transition Path Theory (TPT)

The kinetic property analysis can be performed after completing the estimation and the validation of the MSM. So far, to build the MSM of histone tails, the conformational space is divided into a set of discrete microstates, and the microstates are grouped into similar conformations to get macrostates. Now, the MSM network can be obtained based on the transition *n x n* probabilities to go from one state to another at a specific lag time estimated based on the implied timescales. Each histone tail has about four to five macrostates; therefore, the transition probabilities between these macrostates have been computed to build MSM network plots. We have obtained transition probabilities between the histone tails macrostates for the WT H3 tails, H4 tails, H2A tails, H2B tails, and acetylated H2B tails. These probabilities are plotted on the MSM network plot with transition probability shown next to the arrow to indicate the transition from one state to another. **(*SI Appendix* Fig. S11 – S15)**.

After computing the transition probabilities, we also calculate the mean first passage time (MFPT), as the MFPT provides the mean transition time between the states of the histone tails in the MSM. The MFPT from the initial state *i* to the final state *j* can be determined by this equation *MFPT*_*if*_ = ∑_*j*_ *T*(τ)_*ij*_ (τ + *MFPT*_*if*_). Here, τ is the lag time, and T(τ) is the corresponding transition probability matrix. To calculate the MFPT from the microstate MSM, the microstates within the destined macrostates need to set the MFPT as zero. Then, MFPTs from each microstate in the starting macrostate are calculated. A weighted average is obtained as *MFPT*_*if*_. *MFPT*_*if*_ = ∑_*l*∈*i*_ *p*_*i*_*MFPT*_*lf*_ here *p_l_* is the normalized population of the microstate *l* within the macrostate *i.* The kinetic information of the histone tails is extracted from the MSM by calculating the MFPTs between the different macrostates (52, 63). The inverse of the MFPT (1/mfpt) provides the rate between the transition states.

The MFPTs between the transition states of the histone Tails-1 **(Fig. 4-7)** and Tails-2 **(*SI Appendix* Fig. S16-S19)** are shown in the nanosecond range on the MSM network plot. Each histone tail macrostate is labeled with their stationary population percentages. In addition, we have used Transition Path Theory (TPT) to calculate the statistics of the transition pathway between the transition states of the histone tails with their corresponding percentages. This provides information regarding the TPT fluxes of the major pathways that histone tails follow.

**Fig. 4.**
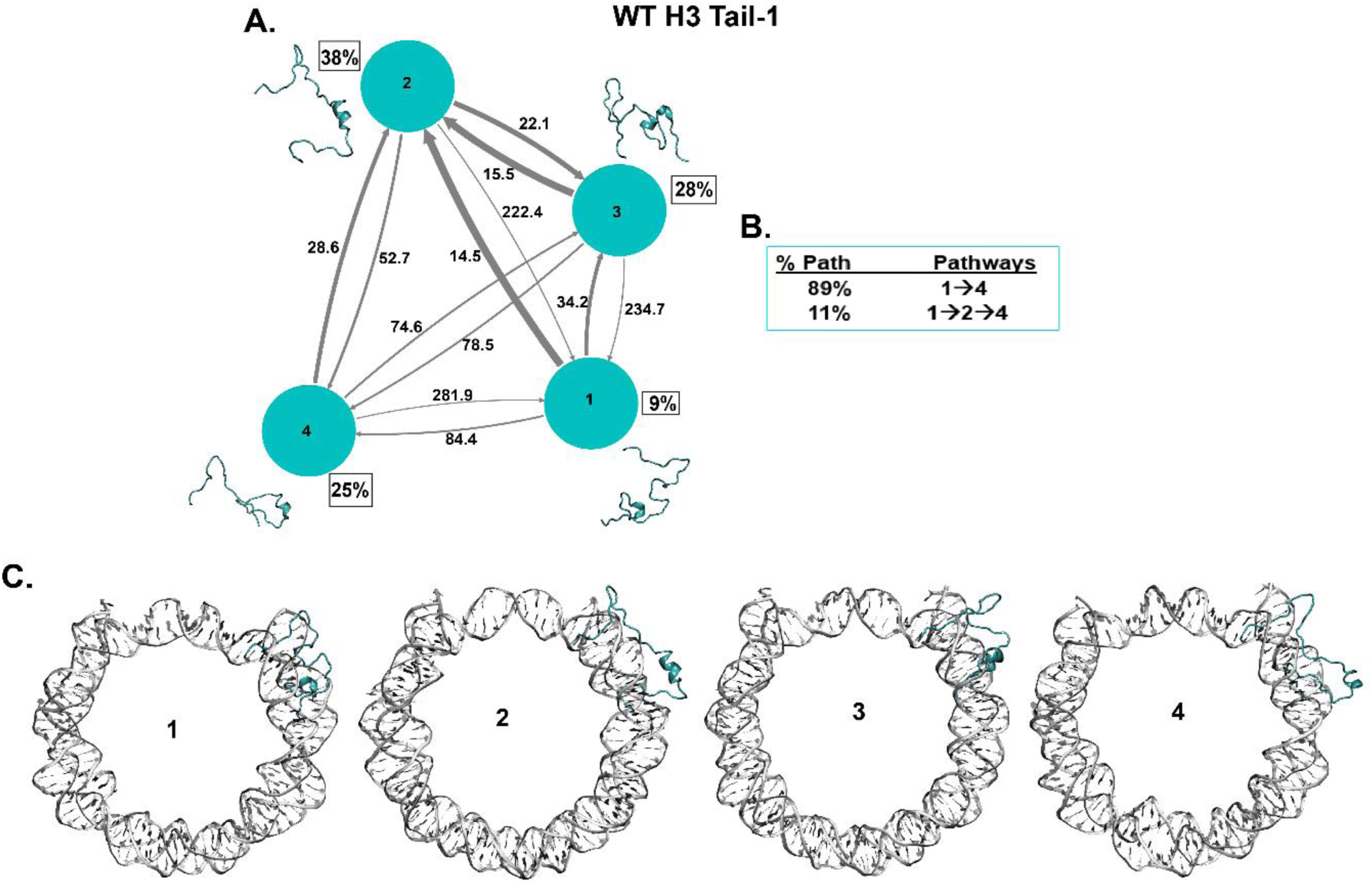
The Kinetic network of conformational states of H3 Tail-1. **(A)** WT H3 Tail-1 (cyan circles) network plots connect four macrostates of H3 tail-1. The corresponding conformations of each states are shown next to each state for H3 tail-1. The population percentages of each conformations are shown next to each state. The macrostates are connected with arrows. The thickness of the arrows is proportional to the rate of the transition and is labelled with its respective MFPT values in nanoseconds. **(B)** The net flux of the network is obtained from Transition Path Theory (TPT) analysis for H3 tail-1. These TPT calculations show major pathways with their path percentages for H3 tail-1. **(C)** The conformational states of H3 tail-1 with NCP DNA is shown. It shows the position of tail (cyan) at each macrostates with respect to DNA (silver).

For the WT H3 Tail-1, the highest population (38%) is state 2, which is a partial helical structure **(Fig. 4A)**, whereas Tail-2 shows the highest population, which is state 1 (37%), less helical compared to Tail-1 **(*SI Appendix* Fig S16A)**. Previous experimental circular dichroism (CD), NMR and MD simulation studies show that the H3 tail does show some helical structures (11, 38, 64, 65). The front part of the tail is slightly disordered for both the H3 tails. It has been seen that H3 tails undergo conformational dynamics at nanoseconds timescales based on NMR and MD simulations (18, 19, 60). The MFPT between all the states for both H3 tails are shown at nanosecond timescales, and from both H3 tails, the MFPT is highest at 506 ns and the lowest one at around 15 ns. The TPT shows two major pathways of both H3 tails, and the pathway from first to last tail conformations 1➔4 accounts for the majority of percentages compared to other pathways **(Fig. 4B, *SI Appendix* FigS16B)**. Now, as the NCP structure is composed of both histone proteins and DNA, it is essential to consider the location of each tail conformational state with respect to the nucleosomal DNA. Our previous MD studies have shown that the histone N-terminal tails can collapse onto the DNA or extend away from the DNA. The tails rapidly fluctuate between these two states. Therefore, when we obtained the transition states of the histone tails based on MSM, we also considered the position of all histone H3, H4, H2A, H2B tails with respect to the DNA, whether it is in a collapsed or extended away from the DNA. Both H3 tails mostly stay collapsed or near the nucleosomal DNA SHL ±7 regions **(Fig. 4C, *SI Appendix* FigS16C)**. However, we have observed that the tail with partial helices structures stay close to DNA, and random coil structures of the tails collapse on the DNA. The collapse of the H3 tails conformations is consistent with previous observations of direct interactions of collapsed H3 tails with the nucleosomal DNA based on NMR single molecule FRET and MD studies (19, 22, 38, 66, 67). Previous MD study from our lab also showed that H3 tails condenses in the minor groove on SHL±7 region and have observed helical conformations of the H3 tails (38). The H3 tails conformation pathways based on the TPT show that tails undergo partial helical to slightly less helical but front of the tail mostly stays collapsed onto nucleosomal DNA.

Further, histone WT H4 Tail-1 shows the highest population (45%) for two states with random coil and turn structures (Fig. 5A), whereas Tail-2 shows the highest population (39%) for state 4, which is a random coil structure **(*SI Appendix* Fig S17A)**. The TPT shows three significant pathways for both H4 tails **(Fig. 5B, *SI Appendix* FigS17B)**. H4 Tail-1 shows 1➔2➔4 as a major pathway, while Tail-2 shows 1➔4 as a major pathway between transition states. The Tail-1 interconverts from random coil to turns and slight helical structure. In addition, the MFPT between different states is shown in nanoseconds timescales, with 15 ns being the lowest and 387 ns being the highest MFPT among both H4 tails. Also, H4 Tail-2 shows predominantly random coil conformations and a slightly faster MFPT than Tail-1. H4 Tail-1 has tail conformations that mostly collapse on the nucleosomal DNA near SHL±2 region; the H4 Tail-2 has random coil conformations that stay collapsed between the two DNA gyres **(Fig. 5C, *SI Appendix* FigS17C)**. Previous NMR and HDX-MS studies show that H4 tails conformational dynamics occur at nanosecond timescales (18, 20, 68). In addition, the H4 tails conformational dynamics of its basic patch interact with adjacent DNA and H2A/H2B acidic patch in nucleosome array (69, 70).

**Fig. 5.**
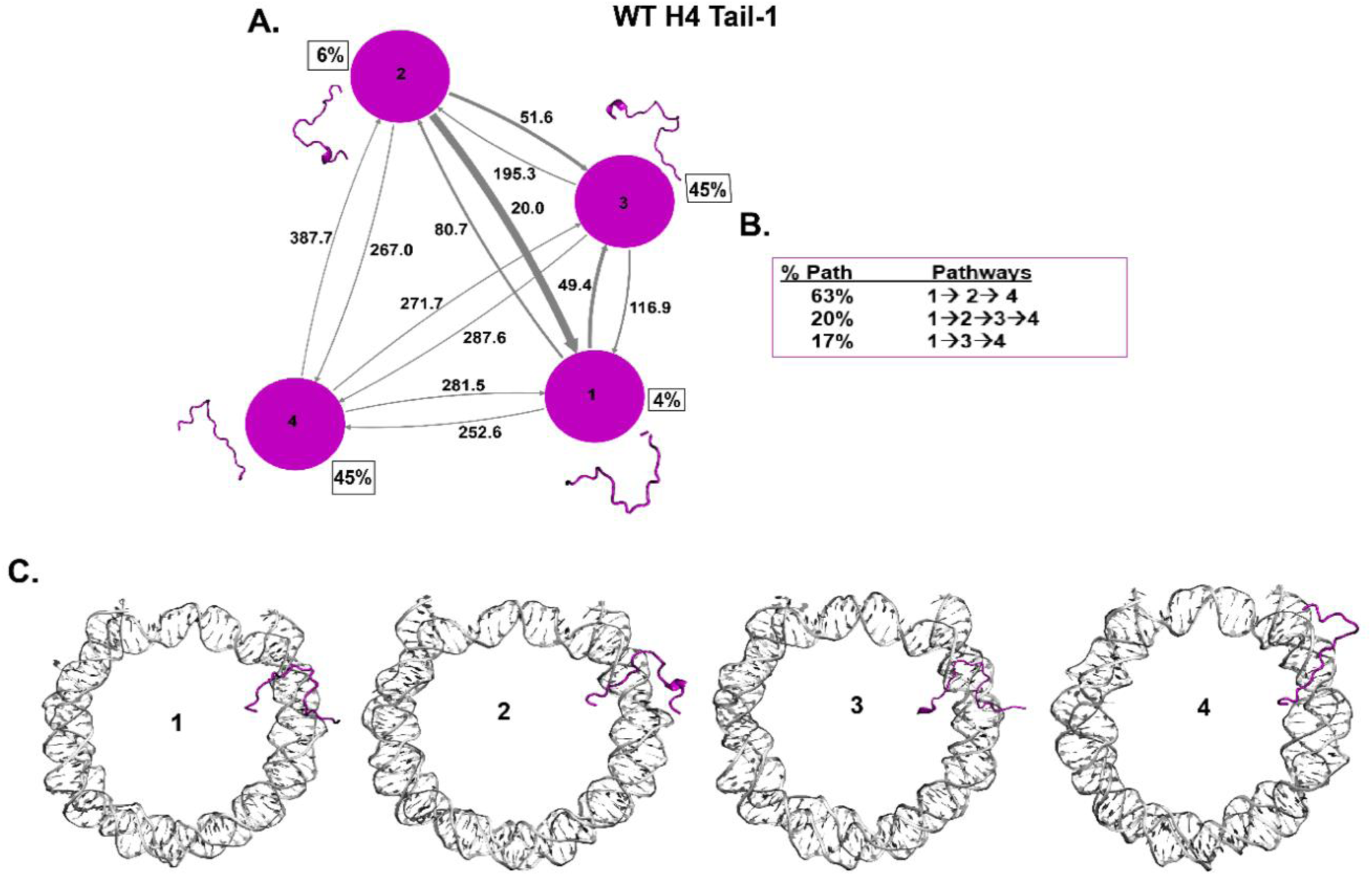
The Kinetic network of conformational states of H4 Tail-1. **(A)** WT H4 Tail-1 (magenta circles) network plots connect four macrostates of H4 tail-1. The corresponding conformations of each states are shown next to each state for H4 tail-1. The population percentages of each conformations are shown next to each state. The macrostates are connected with arrows. The thickness of the arrows is proportional to the rate of the transition and is labelled with its respective MFPT values in nanoseconds. **(B)** The net flux of the network is obtained from Transition Path Theory (TPT) analysis for H4 tail-1. These TPT calculations show major pathways with their path percentages for H4 tail-1. **(C)** The conformational states of H4 tail-1 with NCP DNA is shown. It shows the position of tail (magenta) at each macrostates with respect to DNA (silver).

The histone H2A N-terminal tail is the shortest tail (1-15 residues) among the histone tails of the NCP. Both H2A tails show random coil structures. WT H2A Tail-1 shows the highest population (52%) for state 4, which is a random coil. The other states are also random coils (Fig. 6A). For H2A Tail-2, the highest population (50%) is state 4, which is also a random coil, and other states of Rail-2 are random coils as well **(*SI Appendix* Fig S18A)**. H2A Tail-1 and Tail-2 show major pathways as 1➔4 out of two pathways, and all in-between states are random coils **(Fig. 6B, *SI Appendix* FigS18B)**. The MFPT between different states shows nanosecond timescales with around 5 ns being the lowest and 415 ns being the highest among both H2A tails. Also, NMR and MD simulations study showed that H2A N-terminal tails are unstructured and dynamically faster at nanosecond timescales (11, 20). Further, both H2A tails collapse on the DNA between the two DNA gyres **(Fig. 6C, *SI Appendix* FigS18C)**. The H2A tails collapse around SHL±4 regions and interact with nucleosomal DNA, consistent with the previous studies (71).

**Fig. 6.**
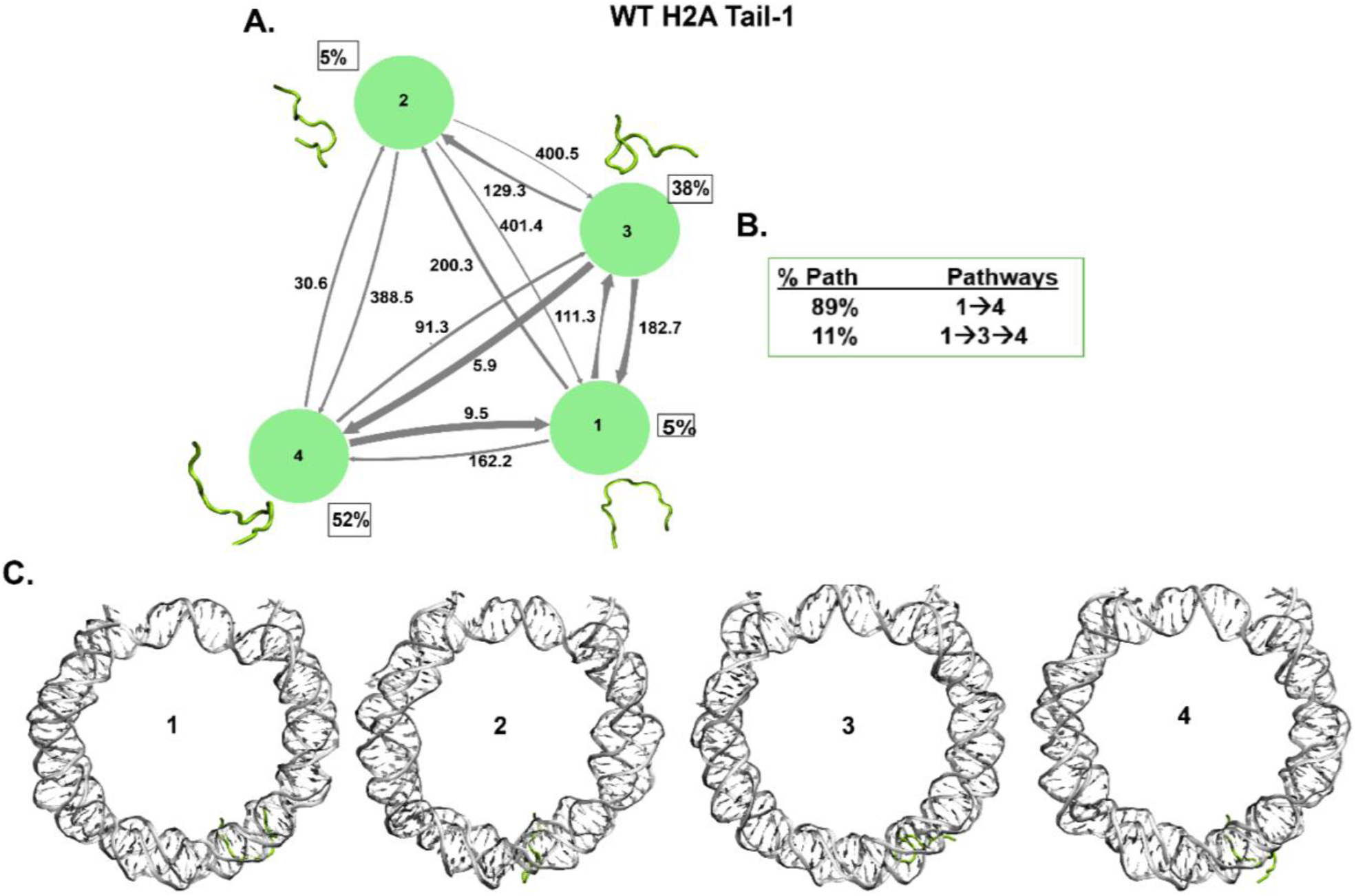
The Kinetic network of conformational states of H2A Tail-1. **(A)** WT H4 Tail-1 (green circles) network plots connect four macrostates of H2A tail-1. The corresponding conformations of each states are shown next to each state for H2A tail-1. The population percentages of each conformations are shown next to each state. The macrostates are connected with arrows. The thickness of the arrows is proportional to the rate of the transition and is labelled with its respective MFPT values in nanoseconds. **(B)** The net flux of the network is obtained from Transition Path Theory (TPT) analysis for H2A tail-1. These TPT calculations show major pathways with their path percentages for H2A tail-1. **(C)** The conformational states of H2A tail-1 with NCP DNA is shown. It shows the position of tail (green) at each macrostates with respect to DNA (silver).

### Effects of H2B N-terminal Acetylation for MSM Analysis

The MFPTs between the transition states of the WT and ACK H2B tail-1 are shown in the nanoseconds range on the MSM network plot **(Fig. 7A-B)**. The highest population (61%) for the WT H2B Tail-1 is state 4, which is a random coil structure, whereas the highest population (35%) for the ACK H2B Tail-1 is state 1, which is the helical structure. Similarly, for H2B Tail-2, the highest population for WT H2B tail-2 is state 4 (56%) and for ACK H2B Tail-2 is state 5 (40%) **(Fig. 7A-B, *SI Appendix* FigS19A-B)**. Both states show random coils. However, the transition states from state 1 to the last state for acetylated H2B Tail-2 involves more helices in the H2B tail than the WT H2B Tail-2. Thus, both ACK H2B Tail-1 and Tail-2 do involve more helix propensity compared to the WT. We have also observed this phenomenon in our previous study, where there is an increase in helicity in the H2B tail upon acetylation (34). In addition, the experimental study by Wang et al (17) has described that lysine acetylation increases the helical content of the histone tails by performing circular dichroism (CD) analysis.

**Fig. 7.**
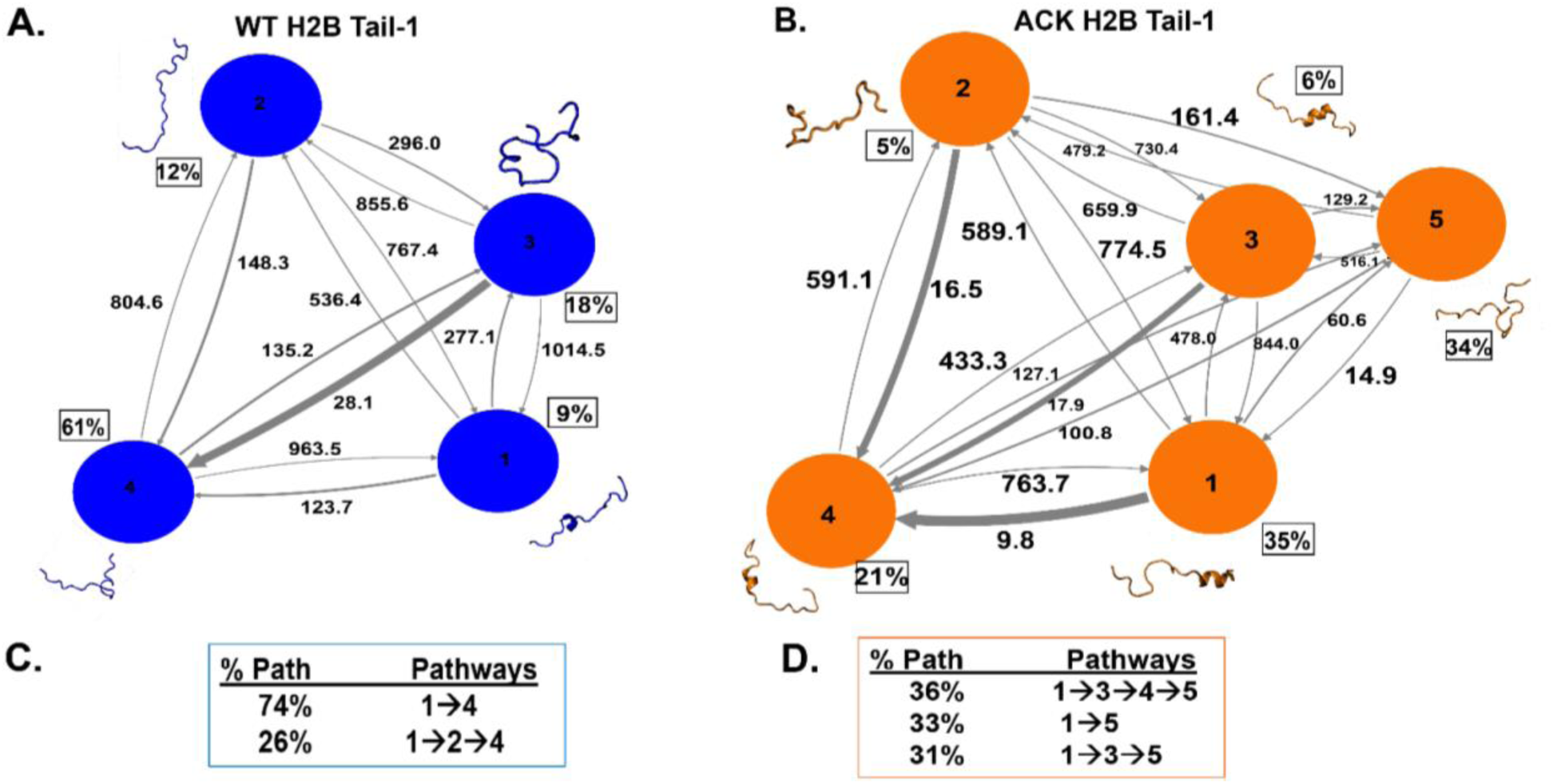
The Kinetic network of conformational states of H2B Tail-1. **(A)** and **(B)** WT (blue circles) and ACK (orange circles) H2B Tail-1 network plots connect four and five macrostates respectively for H2B tail-1. The corresponding conformations of each states are shown next to each state for both WT (blue) and ACK (orange) H2B tail-1. The population percentages of each conformations are shown next to each state. The macrostates are connected with arrows. The thickness of the arrows is proportional to the rate of the transition and is labelled with its respective MFPT values in nanoseconds. **(C)** and **(D)** The net flux of the network is obtained from Transition Path Theory (TPT) analysis for both WT and ACK H2B tail-1. These TPT calculations show major pathways with their path percentages for both WT and ACK H2B tail-1 systems.

Further, we compute the TPT flux between macrostates **(Fig. 7C-D, *SI Appendix* FigS19C-D)**. The TPT calculation shows the major pathways between the transition states of the H2B tails for both the WT and ACK systems with their corresponding percentages. This provides us with information regarding the major pathways that histone tails follow to exert the functions. Earlier, we showed that histone N-terminal tails can collapse onto the DNA or extend away from the DNA during the MD simulations. The tails rapidly fluctuate between these two collapse and extended conditions. In acetylation, the tails mostly extend away from DNA (34). Therefore, when we obtained the transition states of the histone H2B tails for WT and ACK based on MSM, we also considered the position of each conformations, whether they are collapsed or extended away from the DNA for H2B Tail-1 and Tail-2 (**Fig. 8A-B, and *SI Appendix* Fig. S20A-B)** for both the WT and ACK systems. As the acetylated H2B tails have four lysine residues that are acetylated, it shows that three out of five states of the H2B Tail-1 are extending away from the DNA. Most of the interactions of each histone tails bound to the DNA are due to strong electrostatic interactions via salt bridge formation between the positively charged amino acids of the tails and the negatively charged phosphate backbone of the DNA. When the lysine residues of the H2B tails are acetylated, the H2B and DNA interactions decrease due to the charge neutralization of the lysine residues. As the interactions break between the tail and the DNA, the H2B tail extends out and stays away from the DNA. As shown in Fig 8B and Fig S20B, the majority of the acetylated H2B tail conformational states extend out and stay away from the DNA compared to the WT. For example, the acetylated H2B tail-2 conformation state 5, which shows a majority population of 40% (***SI Appendix* Fig. S19B)**, stays away from the DNA **(*SI Appendix* Fig.S20B).** The same H2B tail for the WT, the conformation state 1, which shows the majority population of 56% stays close to the DNA and between the two DNA gyres around SHL±5 and SHL±3 regions. Similarly, for the acetylated H2B Tail-1, the first state with the highest population is helical and extended away from the DNA (**Fig. 7B and Fig. 8B)**.

**Fig. 8.**
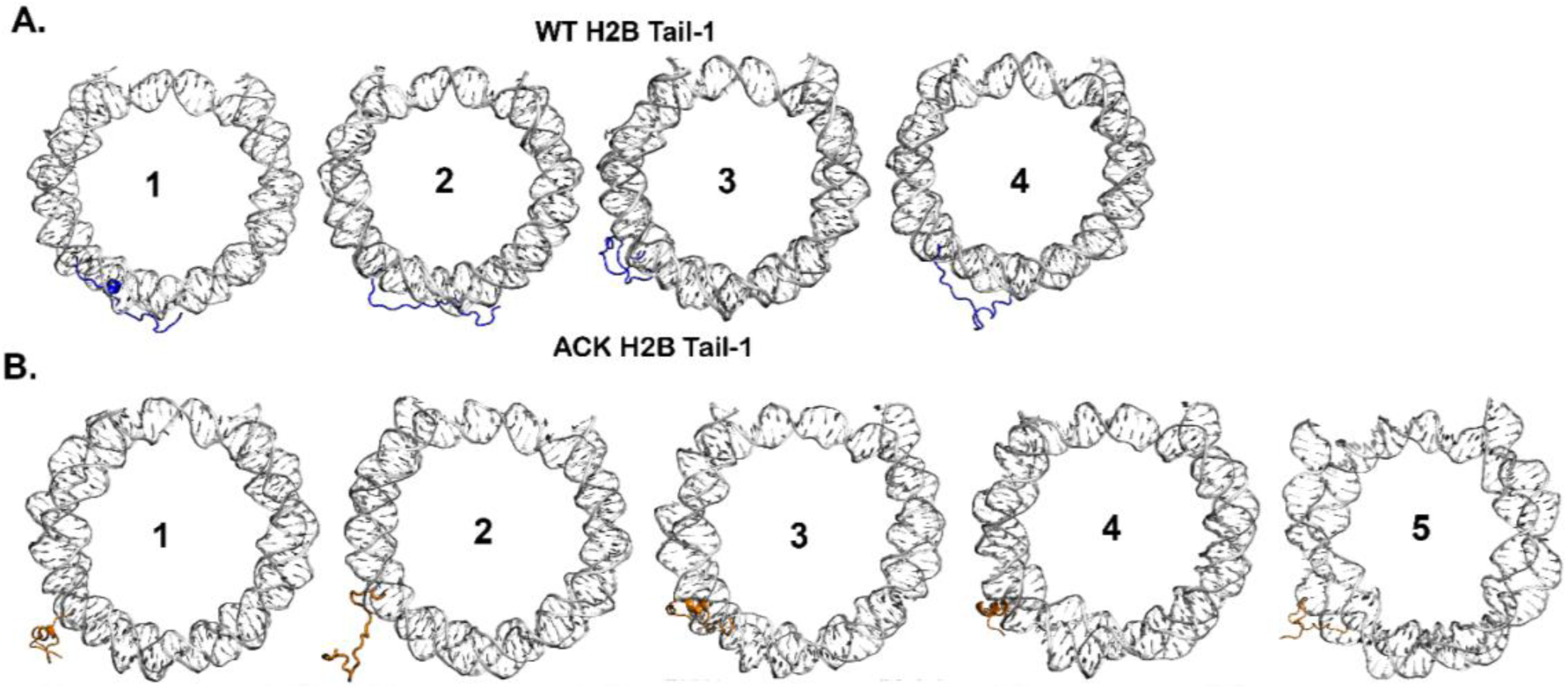
The conformational states of H2B Tail-1 with DNA of NCP. **(A)** and **(B)** WT (blue) and ACK (orange) H2B Tail-1 conformational states that are shown in Figure 4, the same conformational states of the tail with NCP DNA is shown here. It shows the position of each tail conformations at each macrostates shown earlier with regards to the DNA whether the tail collapsed to the DNA or elongated outwards from the DNA.

The kinetic network shows multiple pathways via TPT of WT and ACK H2B Tail-1 and Tail-2 **(Fig. 7C-D, *SI Appendix* Fig. S19C-D).** We have identified the pathway for the WT system for H2B Tail-1 that accounts for 74% (1➔4) of the total pathways, and the majority of these transition states from first to last states of the H2B tails that either collapse onto the DNA or are closer to DNA around SHL±5 and SHL±3. In this major pathway, the H2B Tail-1 for WT that was initially in between two DNA gyres has slightly moved out of two gyres yet stayed in proximity to the DNA. For the acetylated H2B tail-1, there are three pathways, and one of the pathways 1➔5 shows a change in the secondary structure conformation and the tail stays away from the DNA. The intermediate transition states from the first to the last state show tail fluctuation between the collapse of the DNA and extension away from the DNA. As not all positively charged lysine residues of the H2B tails are acetylated, other positively charged residues still attempt to interact with DNA. For WT H2B Tail-2, the major pathway that accounts for 84% (1➔4) of the total pathways, shows the tail location that is present between DNA gyres shifts in closer proximity to the DNA. The ACK H2B Tail-2 is the major pathway 1➔3➔5, the tail conformation shifts from helices to random coil and stays away from the DNA. As the DNA has SHL regions from SHL 0 to SHL±7, the H2B tails mainly interacts with the SHL± 5 and SHL± 3 regions. Therefore, when the tail conformations collapse onto the DNA or have stayed near the DNA, they mostly interact with the SHL± 5 and SHL± 3 regions of the DNA.

## Discussion

Herein, we aim to elucidate the kinetic properties of the conformational dynamics of histone tails H3, H4, H2A, H2B, and acetylated (ACK) H2B that can provide insights into their biological functions. We perform a total of 12 µs long all-atomistic MD simulations of two NCP systems (PDB: 1KX5) at 0.15M salt concentration of unacetylated (WT) and H2B tail lysine-acetylated (ACK) systems. The acetylated lysine residues for both H2B tails are Lys 5, 12, 15, and 20 at 0.15 M salt concentration. These particular lysine residues are associated with the tumor repressor protein p14ARF (72) and ATF2 (73) coactivator. Both p14ARF and ATF2 proteins maintain transcription through H2B tail interactions through acetylation during gene regulation. The interaction of p14ARF with H2B tails involves deacetylation via HDAC1, leading to transcription repression, and in the absence of p14ARF, HAT acetylates these four lysine residues, leading to transcription activation. The p14ARF controls apoptosis or cell death in response to oncogenic stress and regulates gene transcription; however, p14ARF is often mutated in many human cancers. In addition, ATF2 (activating transcription factor 2) has been associated with H2B and H4 acetylation. It is a coactivator for p300 HAT, but ATF2 also has HAT activity that acetylated H2B tail lysine at positions 5, 12, and 15, which results in transcription activation. The H2B N-terminal tail is a critical regulator in gene expression and is ubiquitous in diseases, yet the dynamics and kinetics of the H2B tails remain to be understood.

Histone N-terminal tails are disordered, yet they undergo transient secondary structure conformation changes. Our previous study, Patel *et* al. (34) has shown that the H2B histone tails undergo conformational changes by transitioning from one state to another throughout MD simulations. Using principal component analysis (PCA) of the H2B N-terminal tails for WT and ACK systems, we show distinct free energy basins belonging to specific H2B tail conformations. The dynamic properties of histone tails, such as the kinetics of transitions among different conformations need to be further elucidated. There is one computational study by Zheng et al. (63) that focuses on the H3 tail free energy landscape and underlying kinetics, but this study does not consider the entire NCP in building their MSM. Overall, they use input features such as dihedral angles and show that the H3 tail rapidly transitions between different states with their MFPTs (63). It is essential to consider the tails in complex with the DNA of the NCP to thoroughly understand the tail dynamics. Here, we use Markov state models (MSM) to gain further understanding of the dynamics of histone tails H3, H4, H2A, H2B and ACK H2B transitioning from one state to another in the full NCP complex. A summary of the major findings for each histone tail are in **Table 1**.

**Table 1.** Summary of Histone Tails conformational dynamics and kinetics analysis.

| Histone Tails | Highest Population |  | Structural Conformation | MFPT (ns) |  | Total No. of states | Major Transition Path |
| --- | --- | --- | --- | --- | --- | --- | --- |
|  | State | % |  | Low | High |  |  |
| WT H3 | 2 | 38 % | Partial Helical | 15 | 506 | 4 | 1→4 |
| WT H4 | 3 | 45% | Minor helix/ turn | 15 | 387 | 4 | 1→2→4 |
| WT H2A | 4 | 52% | Random coil | 6 | 415 | 4 | 1→4 |
| WT H2B | 4 | 61% | Random coil | 18 | 1014 | 4 | 1→4 |
| ACK H2B | 1 | 35% | Helical | 4 | 763 | 5 | 1→3→4→5 |

We observe that histone H3 tails and H2B tails show slightly more helical structure of the tails compared to the H2A and H4 tails based on our MSM analysis. H3 and H2B tails have approximately 30% hydrophobic residues, whereas H2A and H4 tails have only about 15%. Helical or globular proteins contain about 50% hydrophobic residues such as phenylalanine, leucine, and isoleucine with bulky hydrophobic groups (74). The H3 tails are mostly collapsed onto the DNA near the SHL±7 regions. Collapse leads to interactions with the DNA and the NCP core regions. These contacts are primarily driven by the positively charged lysine and arginine residues of the H3 tails (19, 22, 38, 66, 67). Our previous MD study has shown that when H3 tails collapse onto DNA, it leads to the outward movement of the SHL±7 region. This results in the unwrapping of the DNA entry/exit region, initiating breathing motion. Upon collapse, the helical content of the tail increases (38). Conformational changes of the H3 tail could facilitate binding to the PHF1 Tudor domain, promoting partial unwrapping of the DNA. This indicates that the H3 tail can regulate DNA binding dynamics (75).

H4 tails conformations are slightly less helical than H3 tails, but like H3 tails, H4 tails also mostly collapse onto the DNA around the SHL±2 region. One of the H4 tails shows a slightly shorter MFPT than the other H4 tail. It has been suggested that there is cross-talk between H3 and H4 tails, which might be arising from changes in the conformational ensembles of one tail’s DNA collapsed/bound state when the other tail is perturbed (14, 76). H4 tails interchange their conformations between random coil to slightly helical. As mentioned, conformational ensembles are part of crosstalk; the kinetics of tail conformations can provide valuable insights into the mechanisms of their interaction and dynamics. H4 tail residues 16-23 include mostly positively charged lysine and arginine, known as the H4 basic patch located close to the NCP core. The conformational dynamics of the H4 tails basic patch forms intra and/or inter-nucleosome interactions with adjacent DNA or the H2A/H2B acidic patch in a nucleosome array (69, 70, 77, 78)

H2A tails are the shortest compared to the other histone tails. This could potentially lead to an increase in their fluctuations with a lower MFPT. Among all the WT histone tails, H3, H4, H2A and H2B, the WT H2A tails show the lowest MFPT. H2A tails exhibit predominantly random coil conformations of the tail. The tails are also collapsed around the SHL±4 regions. Just like how the H3 tail’s collapse onto the DNA contributes to unwrapping, H2A tails are also linked to promoting DNA unwrapping (23, 67). Certain histone tails may influence the dynamics of the other tails. In particular, the H2A N-terminal tails are positioned close to the H2B tails and might influence each other. Notably, the correlation of the H3 and H2A tails can modulate DNA breathing dynamics (14).

Based on the network plot of WT and ACK H2B tails, acetylated ACK H2B tails mostly have alpha helix conformations compared to the other WT tails. Computational and experimental studies have observed the phenomenon that acetylation increases the helicity in the histone tails (17, 34, 79). The transition states between WT H2B tails are predominantly random coils compared to ACK tails. This might be due to the H2B tails being highly positively charged with two Arg and eight Lys residues, which generally creates a repulsive electrostatic interaction that makes it harder to form distinct compact helical structures. Upon acetylation, the repulsion decreases in the tail as there is charge neutralization of the acetylated lysine residues. Adding the bulky acetyl group would create less repulsion and more hydrophobicity, shifting the tail into more compact helical structures. Thus, we observe more helicity of the tail in the ACK H2B tail-1 and tail-2 compared to WT.

Epigenetic modifications such as acetylation in chromatin regulate vital cellular processes such as transcription, DNA damage repair, and gene regulation. Epigenetic modifications are closely linked to diseases such as cancer, neurological disorders, and inflammatory diseases. Acetylation is closely linked to transcription as it makes chromatin transition from tightly packed to loosely packed states, making the NCP nucleosomal DNA more accessible to RNA polymerase II, transcription factors, and other proteins required for gene regulation. It can also help prevent specific proteins from interacting and causing DNA damage, and it is involved in DNA repair. Acetylation of H2B tails leads to faster transition rates and more dynamic tails; this was also observed in a previous NMR study. Our MFPT transition rates are in nanoseconds timescales, which is rapid for acetylation. This agrees with the NMR study that observes the tail conformational changes at the rapid timescales of picoseconds to nanoseconds upon acetylation (18). Although it is often difficult to study tail dynamics using biophysical techniques, one of the solid-state NMR studies has shown that acetylation during PTMs of histone tails leads to an increase in tail dynamics and, subsequently, stronger interactions with other proteins (18, 22). Furthermore, the rapid transitions in the acetylated tail may regulate access of key proteins to the nucleosomal DNA. Previous NMR studies have shown that acetylation increases enzyme activity as DNA becomes more accessible for enzymes to conduct biological functions. For example, acetylation of histone tails, specifically H2B and H2A, causes an increase in ligation with LIG3 (DNA Ligase 3) (18). And higher fluctuations for the H2B acetylated tails.

Further, H2B tails in the ACK system have shown compact helical structures upon acetylation. This could serve as a docking site for other proteins that recognize the acetylated sites. Bromodomains (BRDs) are one of the proteins that recognize multiple acetylated lysine residues at the site of the histone tails. As histone tails become more hydrophobic in nature, the tails can induce fit perfectly in the hydrophobic binding pocket of BRD. The BRD is associated with epigenetic reader modules and has been implicated in drug treatment for cancer diseases (29, 80-82). Most of the conformational populations for the H2B acetylated tails are helical conformations compared to WT based on our MSM. We also compute transition path theory (TPT) for transition state pathways for the WT and ACK H2B tails. The histone tails fluctuate between collapsing onto the DNA near SHL±3 and SHL±5 regions and extending away from the nucleosomal DNA. The tail conformations of the macrostate in the ACK system mostly show the tail extending away from the DNA. The tail conformations of the ACK H2B tails that stay in proximity to DNA in helical structures can provide docking to other proteins to access the nucleosomal DNA of the NCP to carry out biological functions. Other histone tails, H3, H4, and H2A, show tail conformations mostly collapsed onto the DNA or are present between the two DNA gyres.

The histone tails have been implicated in assembling higher-order chromatin structures including inter-nucleosome interactions, nucleosome array compaction and array oligomerization. H4 tail interactions with an acidic patch of the adjacent nucleosomes in a tetra-nucleosome show that H4 tail interacting with the acidic patch and at the interface between tetra -nucleosomes form the stable nucleosome array structure. This also provides the importance of tail collapse to the DNA making DNA-tail contacts. In addition, previous MD studies show that all the tails form inter-nucleosome contacts primarily through DNA-tail interactions (14, 78, 83). Also, when the tail is modified through PTM, such as acetylation, the conformational ensemble of the tail becomes more compact, promoting intra-nucleosomal interactions rather than inter-nucleosomal interactions (29, 84). As we observed, H2B acetylation shows faster MFPT; this can facilitate a quick transition from inter- to intra-nucleosome interactions in the regulation of higher-order chromatin structure. The faster transition between states can also facilitate quick searching for new binding proteins as the tail decreases its interactions with nucleosomal DNA (14).

As we have observed, histone tails alter their conformations, and upon acetylation, tails alter their conformations compared to WT. These conformations may also contribute to crosstalk. The conformational ensemble of one tail may alter the adjacent tail. As H2B and H2A tails are closer, the changes to the H2A tail may alter the H2B conformations. Both tails interact with DNA SHL regions that are closer, so the changes to one tail’s DNA interactions may alter neighboring tail. A recent study found that acetylation of the H4 tail can affect the conformational dynamics of the H3 tail (14, 21). The conformations of the tails are likely a critical contributor to chromatin PTM crosstalk. Therefore, studying the kinetics of the tail conformations can provide insight into crosstalk.

In conclusion, we perform long-time all-atomistic molecular dynamics simulations of the NCP at 0.15 M NaCl concentration for WT (unacetylated) and ACK (acetylated H2B tail) systems. The four lysine residues K4, K12, K15, and K20 of both H2B tails are acetylated. As histone tails are dynamic, the kinetics of these tails needed to be elucidated. Here, we analyze the kinetics of the histone H3, H4, H2A, H2B tails, and ACK H2B tails to assess the structural dynamics of the histone tails. Our results highlight that acetylated H2B tails show faster transition rates when transitioning from one state to another based on their mean first passage times (MFPT) compared to WT systems. This aligns well with a previous NMR study that observed a similar phenomenon. H2B tail acetylation leads to the tail’s structural changes, making it more helical, which can serve as docking sites for other proteins. Acetylated H2B tails also show more fluctuations in tail motion compared to WT; rapid fluctuations of the ACK tails support regulatory protein activity for biological functions and nucleosome plasticity.

## Materials and Methods

### Simulation Methods

The nucleosome core particle (NCP) was simulated in a physiological salt concentration of 0.15 M NaCl. The initial structure configuration of NCP was obtained from the X-ray crystal structure.(37) as reported in the Protein Data Bank (PDB ID: 1KX5). Both subunits of H2B histone proteins N-terminal tails at K5, K12, K15, and K20 positions of 1KX5 NCP were acetylated by adding an acetyl group using PyMol. The acetylated (ACK) and wild-type (WT) unacetylated H2B tails were parameterized using AMBER forcefields.

Histone proteins of NCP were parameterized with ff19SB(85), and DNA was parameterized using OL15(86). The OPC water model(87) was used, with its Lennard-Jones interaction (Na^+^/OW) modification, using the Kulkarni *et al*. method that provides better estimates of the osmotic pressure(88). For sodium (Na^+^) and chlorine (Cl^-^) ions, Joung and Cheetham(89) parameters were used. Mg^+2^ modification was performed using the Li *et al* parameter method (90). All the forcefields were sourced using the tleap module of AmberTools21 to create the topology and coordinate files for the initial ACK and WT systems. All systems were initially minimized and equilibrated for 100 ns, followed by production runs total of 12 µs. The production runs were carried out using Anton2(91) supercomputer.

The N-terminal residues for tails: H3 (residues 1-43), H4 (residues 1-23), and H2A (residues 1-15) and H2B (residues 4-30). We used PyEMMA(62, 92) version 2.5.7 to construct the Markov State model (MSM) for all the histone tails. Details of MD simulations and MSM construction are provided in SI Appendix.

## Acknowledgments

This work was supported by grants from the NIH (1R15GM146228-01). R. P. is grateful for support from The Rosemary O’Halloran Scholarship. Anton 2 computer time was provided by the Pittsburgh Supercomputing Center (PSC) through Grant R01GM116961 from the National Institutes of Health. The Anton 2 machine at PSC was generously made available by D.E. Shaw Research. We thank Prof. Vincent Voelz for discussions.

## Data Availability Statement

Analysis codes are available on GitHub Repository. https://github.com/CUNY-CSI-Loverde-Laboratory/Histone_N_terminal_Tails_Conformational_Dynamics_Kinetics

Trajectories are available on Zenodo.

