## Supplementary Information for "Unveiling the Conformational Dynamics of the Histone Tails Using Markov State Modeling"

*Sharon M. Loverde

PNAS strongly encourages authors to supply an ORCID identifier for each author. Do not include ORCIDs in the manuscript file; individual authors must link their ORCID account to their PNAS account at www.pnascentral.org. For proper authentication, authors must provide their ORCID at submission and are not permitted to add ORCIDs on proofs.

**Author Contributions:** R.P parameterized and conducted molecular dynamics simulations, analyzed data, prepared all the figures, and wrote the paper. The study and paper writing are supported by S.M.L. S.M.L also supervised the project.

**Competing Interest Statement:** None.

**Classification:** Biophysics and Computational Biology, Physical Science.

**Keywords:** Nucleosome, Markov state Model (MSM), Histone tails, Molecular Dynamics simulations.

**This PDF file includes:**

Figures S1 to S20
Tables S1

### Materials and Methods

### Simulation Methods

The nucleosome core particle (NCP) was simulated in a physiological salt concentration of 0.15 M NaCl. The initial structure configuration of NCP was obtained from the X-ray crystal structure(1) as reported in the Protein Data Bank (PDB ID: 1KX5). Both subunits of H2B histone proteins N-terminal tails at K5, K12, K15, and K20 positions of 1KX5 NCP were acetylated by adding acetyl group using PyMol. The acetylated (ACK) and wild type (WT) unacetylated H2B tails were parameterized using AMBER forcefields. Histone proteins of NCP were parameterized with ff19SB(2), and DNA was parameterized using OL15(3). The OPC water model(4) was used, with its Lennard-Jones interaction ($Na^+$/OW) modification, using the Kulkarni *et al*. method that provides better estimates of the osmotic pressure(5). For sodium ($Na^+$) and chlorine ($Cl^-$) ions, Joung and Cheetham(6) parameters were used. $Mg^{+2}$ modification was performed using the Li *et al* parameter method(7). All the forcefields were sourced using the tleap module of AmberTools21 to create the topology and coordinate files for the initial ACK and WT systems. The number of total molecules and water/ions are shown in Table S1.

All systems were initially minimized and equilibrated for 100 ns, followed by production runs total of 12 µs. The production runs were carried out using Anton2(8) supercomputer. The minimization was done to reduce unfavorable stress by using conjugate gradient and steepest descent gradient for 40 ps. Following minimization, the heating was performed by increasing the temperature of the system to 310 K under *NVT* conditions. Afterward, the systems were equilibrated to 100 ns under *NPT* conditions. The Langevin(9) dynamics method with the collision frequency with a friction constant of 1 $ps^{-1}$ was used to control the temperature of the system. The pressure of the system was controlled by the Berendsen(10) barostat. The simulation was continued for production runs under *NPT* conditions with a 2 fs each timestep. All simulations used the SHAKE(11) algorithm to constrain the bonds involving hydrogen. The Lennard-Jones cutoff value for nonbonded interactions was 12 Å, and electrostatic interactions were treated with the Particle Mesh Ewald (PME)(12) method with full periodic boundary conditions.

### Markov State Model (MSM) construction

Markov state Models (MSMs) have been a well-known tool in protein folding, ligand binding and conformational dynamics. To build Markov modeling, the features, such as torsion angles, distances, etc which best represent the slow dynamics of a system are used. Here, we use backbone torsions and pairwise distances of histone N-terminal tails as input features as we observe that tails undergo secondary structure rearrangement. For both WT and ACK systems, the backbone torsion angles and pairwise distances as features are selected for MSM construction. The N-terminal residues for tails: H3 (residues 1-43), H4 (residues 1-23), and H2A (residues 1-15) and H2B (residues 4-30).

We used PyEMMA(13, 14) version 2.5.7 for all trajectories for featurization using the backbone torsions of N-terminal H2B tail residues of WT and ACK systems. Time dependent independent component analysis (TICA) is a commonly used method for dimensionality reduction. To apply TICA, instantaneous (C (0)) and time lagged (C((τ)) covariance matrices is computed with elements $C_{ij}(0) = \langle X_i(t) X_j(t)\rangle_t$ and $C_{ij}(\tau) = \langle X_i(t) X_j(t+\tau)\rangle_t$ where $X_i(t)$ represents $i^{th}$ feature at time t after the mean has been removed(13, 15). Time-lagged independent component analysis (TICA) is computed for histone tails with a lag time (τ) 5 ns. A *k*-means clustering algorithm is used to conformationally cluster the low-dimensional TICA projections to define microstates. The free energy landscape of the first two slowest TICA-dimensions are plotted for both WT and ACK systems. The free energy landscape is defined as using $\Delta G(x,y) = -RT \ln[P(x,y) / P_{max}]$, where P(x,y) is the probability density distribution, R is the gas constant, T is the temperature (310 K), and

$P_{max}$ is the maximum probability. TICA conformational space is segmented into *k*= 200 cluster centers providing 200 microstates of the tails. MSM estimation is done by calculating implied timescales (ITS). The estimation of ITS computed from eigenvalues ($\lambda i$) at each of the given lag time using $ti = -\frac{\tau}{\ln \lambda i}$, where $\tau$ is the lag time(13, 16). Once the lag time is estimated, to check whether a given transition probability matrix P(τ) is approximately Markovian using the CK test. The MSM validation is done using the Chapman-Kolmogorov (CK) test. The CK property of Markovian matrix is $P(k\tau) = P^{k}(\tau)$ where the left-hand side of the equation corresponds to an MSM estimated at lag time $(k\tau)$ and *k* is an integer larger than 1. The right-hand side of the equation is our system's estimated MSM transition probability matrix to the $k^{th}$ power. Based on how well both sides adhere, the MSM is validated. MSM network with a lag time of 5 ns is constructed using microstates. Microstates are assigned to macrostates using the Robust Perron Cluster Cluster Analysis (PCCA+)(17, 18) algorithm. To identify the transition rates between the conformational states, the mean first passage times (MFPTs) are used and the inverse of MFPT provides kinetic rates for each conformational state. In addition, by using PyEMMA(13, 14), we have integrated transition path theory (TPT)(19) to obtain the net transition pathways and their fluxes. Here, TPT analysis provides a flux network from source state A to sink state B that passes through intermediate states. The dominant pathways of the states and their percentages in total are provided for each systems.

### Supplementary Figures

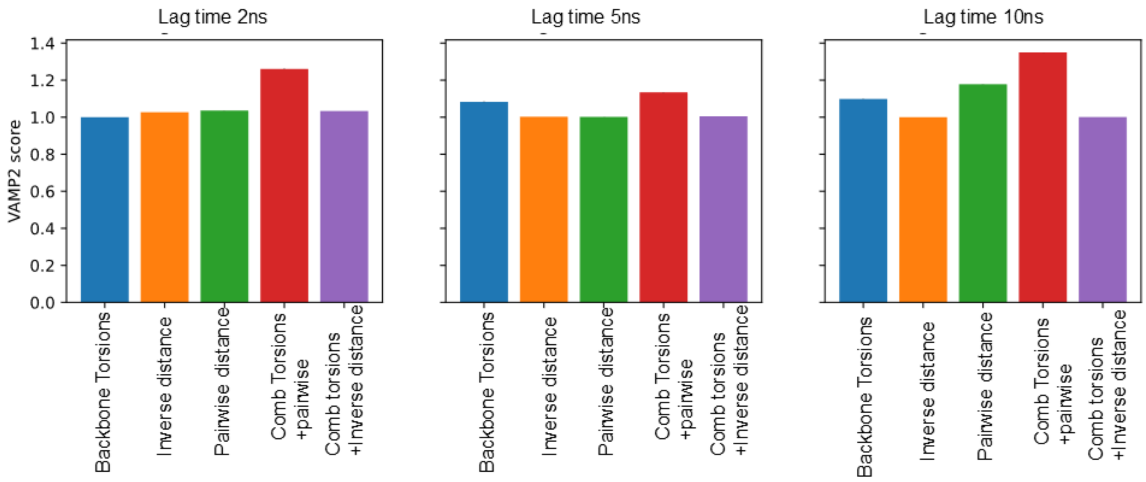


**Fig. S1. Feature selection.** VAMP-2 scores of the various features of histone tails calculated at various lag times. Backbone (blue), inverse distance (orange), $C_\alpha$ pairwise distances (green), combination of the backbone torsions and pairwise distances (red), and combination of backbone torsions and inverse distance (purple). The combination of backbone torsions and pairwise distances is a superior feature across all various lag times and is used for all analysis.

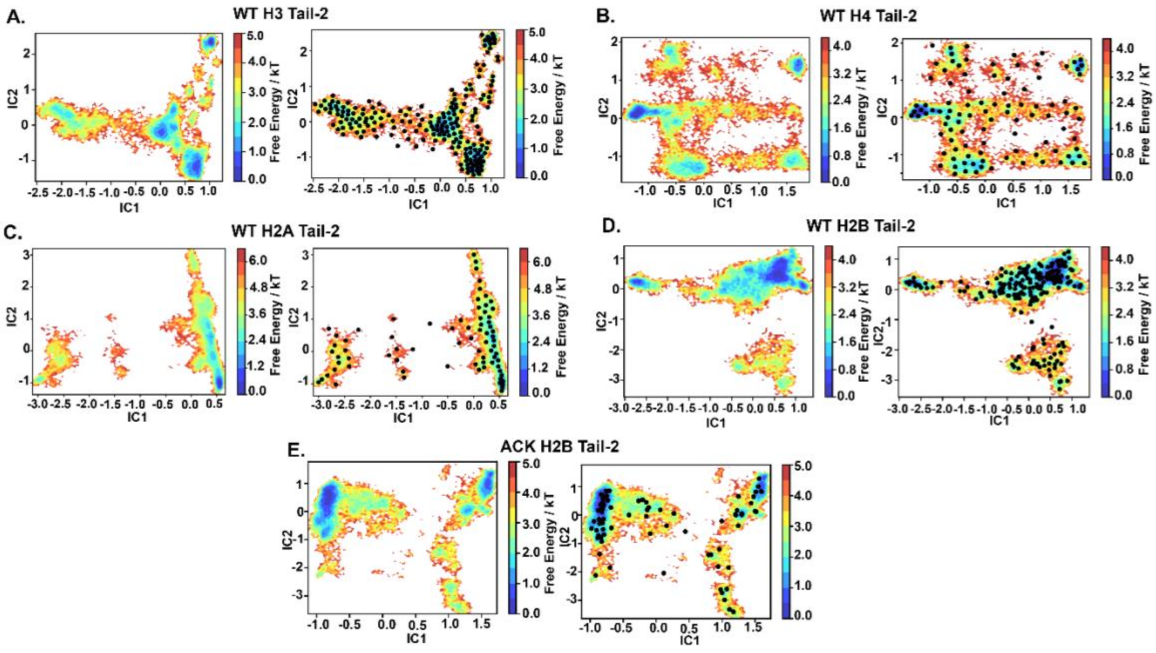


**Fig. S2. Dimensionality Reduction using Time Independent Component Analysis (TICA).** The free energy surface and *k*-means clustering visualizations are projected onto the leading two Independent Components (ICs) for WT **(A)** H3 tail-2 **(B)** H4 tail-2 **(C)** H2A tail-2 **(D)** H2B tail-2 and **(E)** ACK H2B tail-2. The *k*-means clustering provides microstates for each tails based on their conformational space.

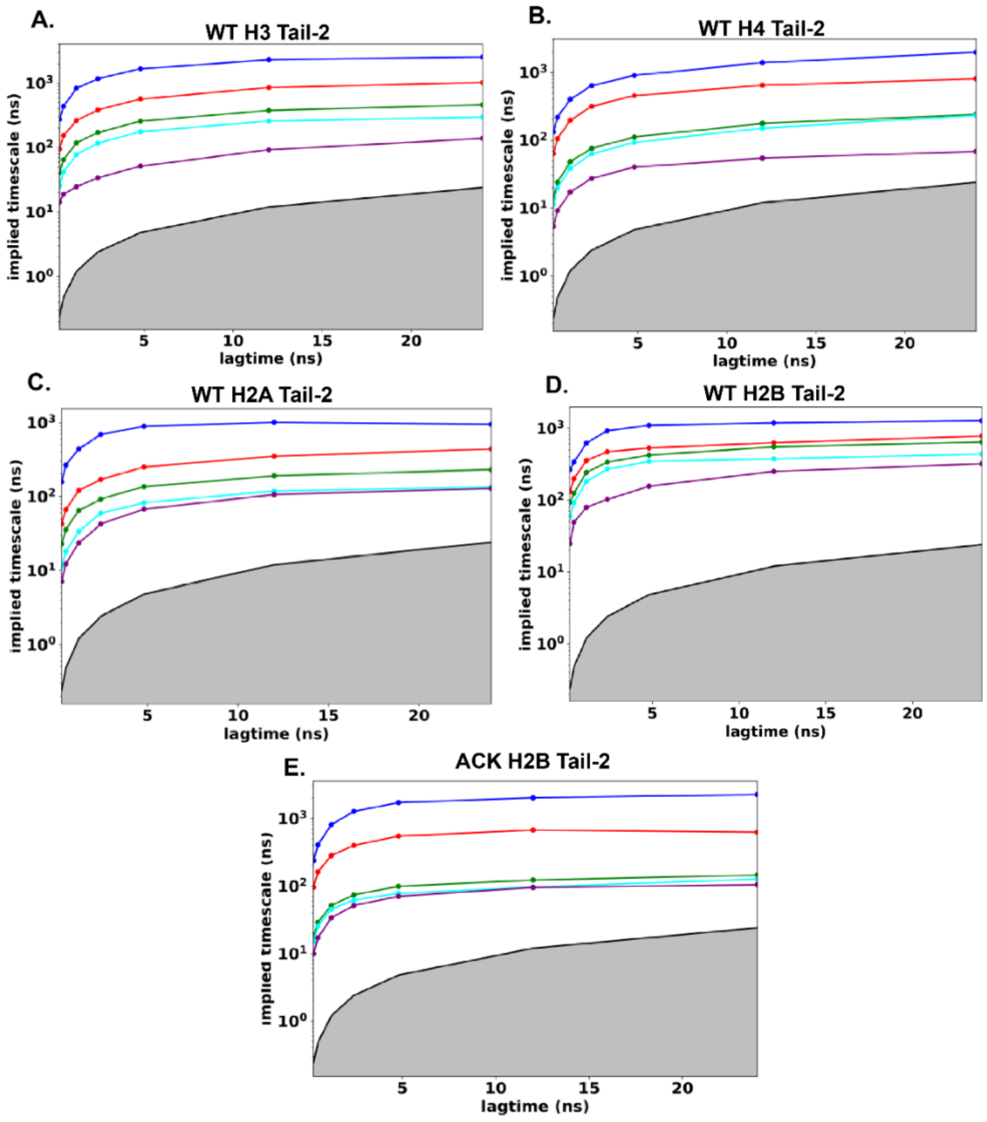


**Fig. S3. Implied Timescales of Histone N-terminal tails.** The implied timescales (ITs) are associated with the five slowest processes for WT **(A)** H3 tail-2 **(B)** H4 tail-2 **(C)** H2A tail-2 **(D)** H2B tail-2 and **(E)** ACK H2B tail-2. The implied timescales plots show Markov processes with different lag times. The solid line corresponds to the maximum likelihood. The blackline with grey shaded area indicates the timescale horizon below which the MSM cannot resolve processes.

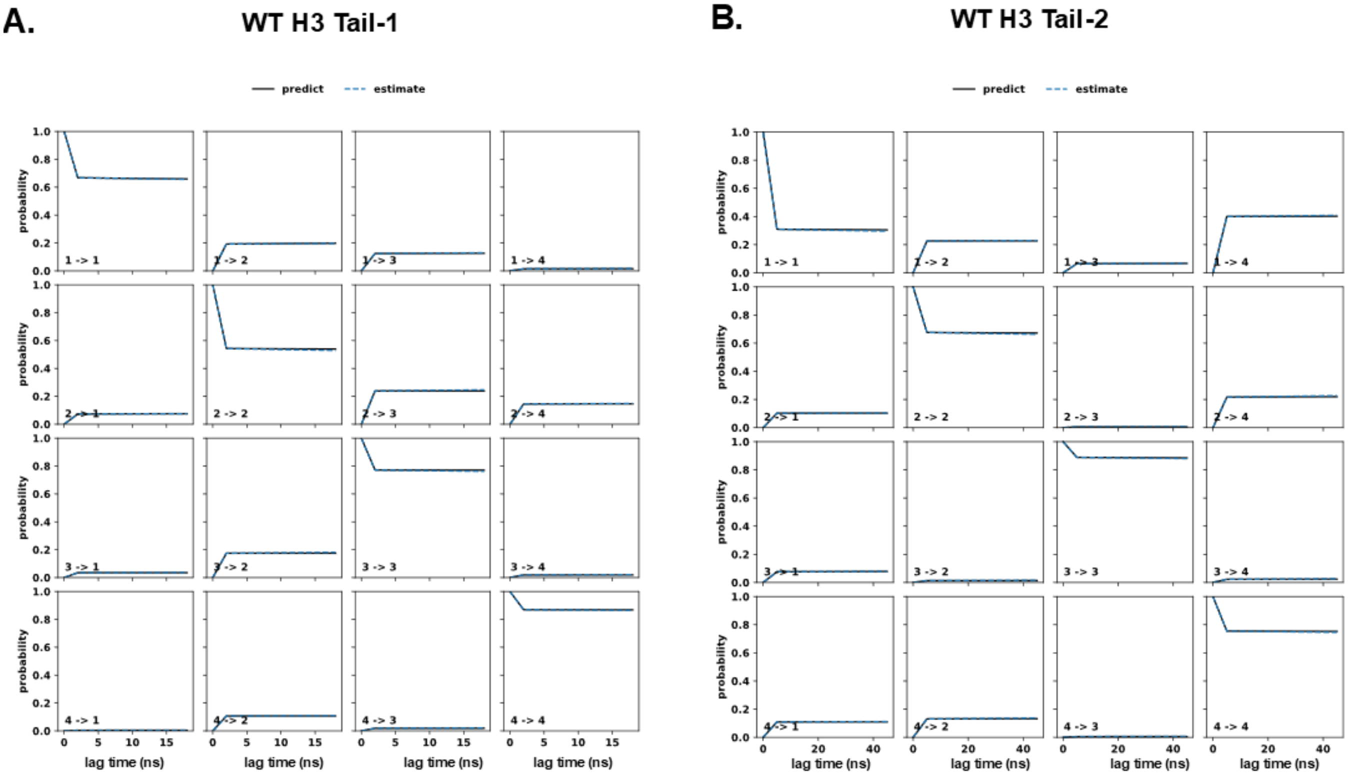


**Fig. S4. Chapman-Kolmogorov (CK) test for MSM validation of H3 Tails. (A)** H3 tail-1 and **(B)** H3 tail-2 show four metastable states. The predictions from our MSM (blue-dash line) agree well with MSM estimated (solid line) for all states.

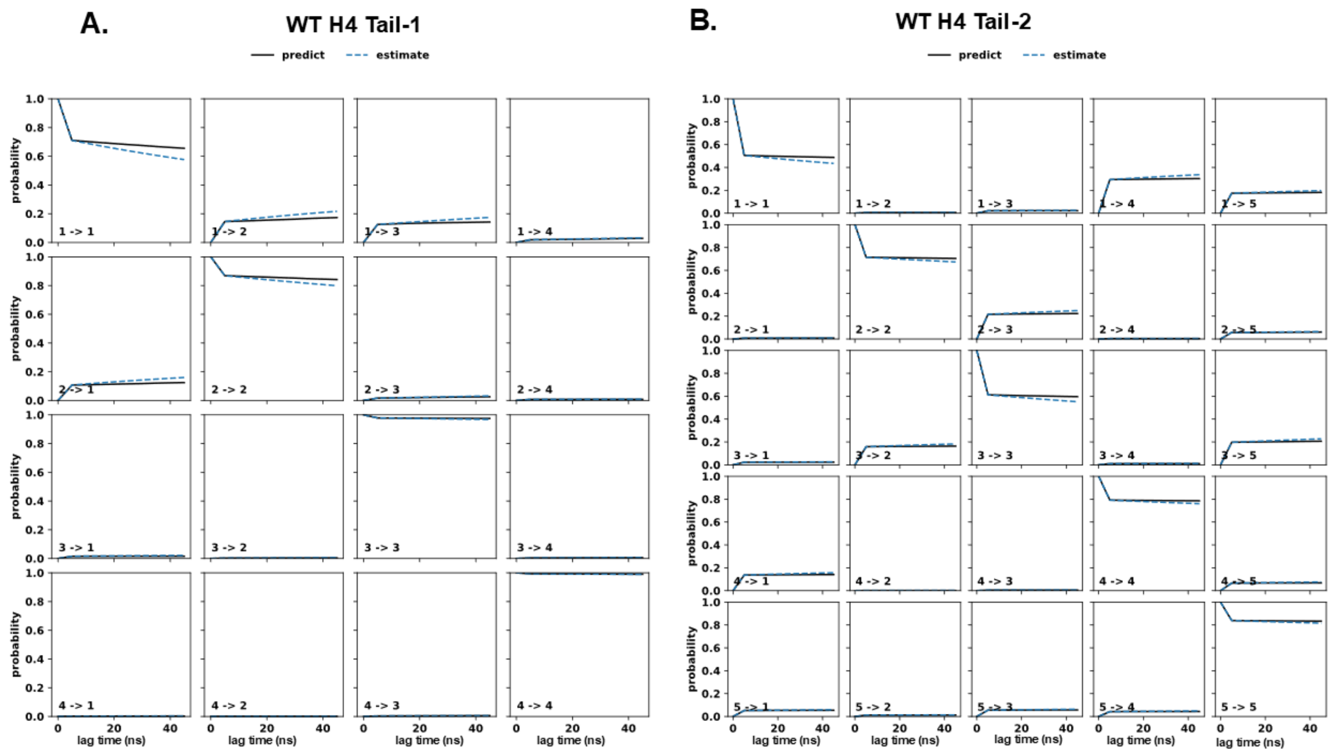


**Fig. S5. Chapman-Kolmogorov (CK) test for MSM validation of H4 Tails. (A)** H4 tail-1 and **(B)** H4 tail-2 show four and five metastable states respectively. The predictions from our MSM (blue-dash line) agree well with MSM estimated (solid line) for all states.

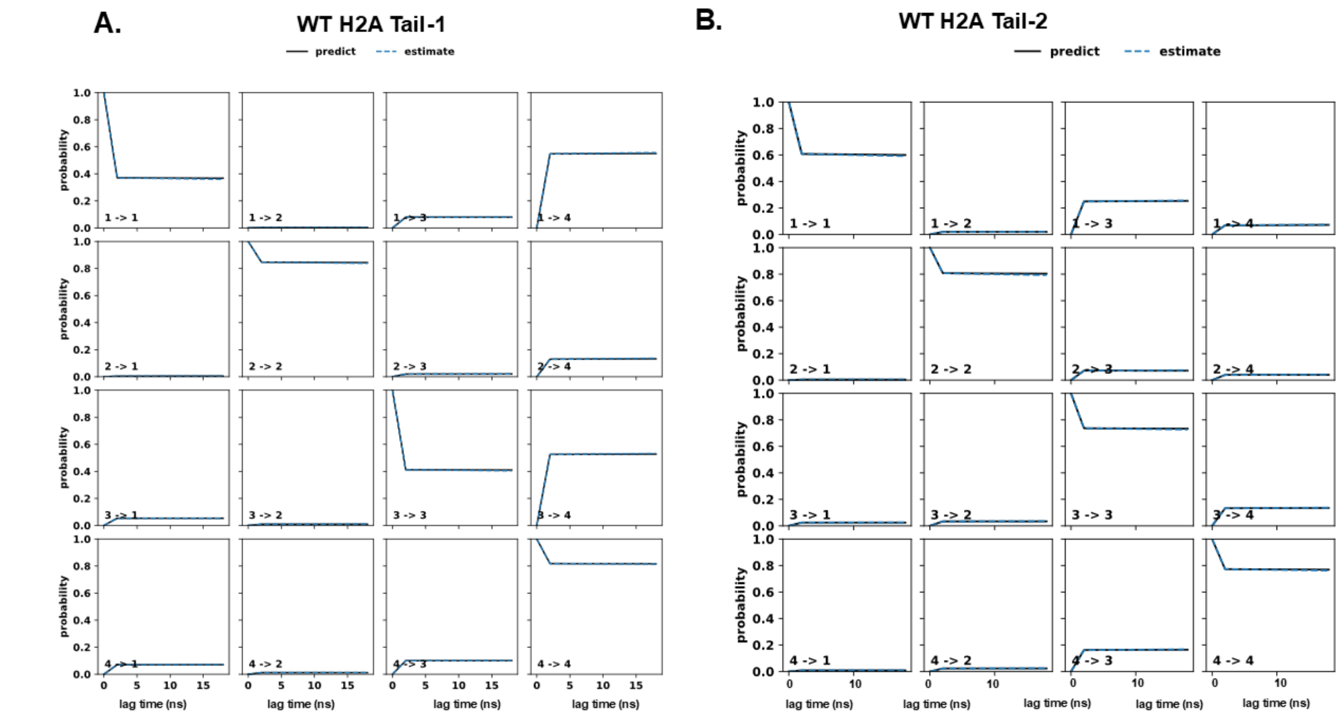


**Fig. S6. Chapman-Kolmogorov (CK) test for MSM validation of H2A Tails. (A)** H2A tail-1 and **(B)** H2A tail-2 show four metastable states. The predictions from our MSM (blue-dash line) agree well with MSM estimated (solid line) for all states.

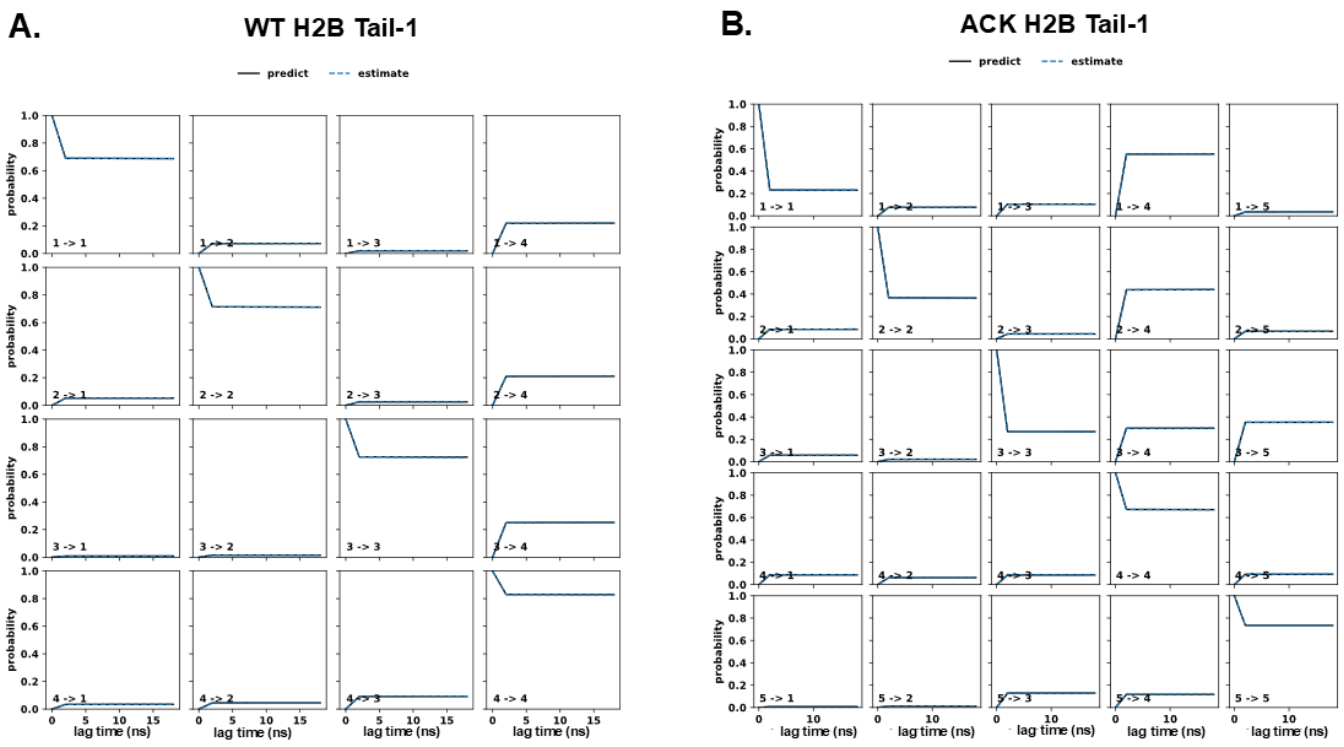

**Fig. S7. Chapman-Kolmogorov (CK) test for MSM validation of H2B Tails-1. (A)** WT H2B tail-1 and **(B)** ACK H2B tail-1 show four and five metastable states respectively. The predictions from our MSM (blue-dash line) agree well with MSM estimated (solid line) for all states.

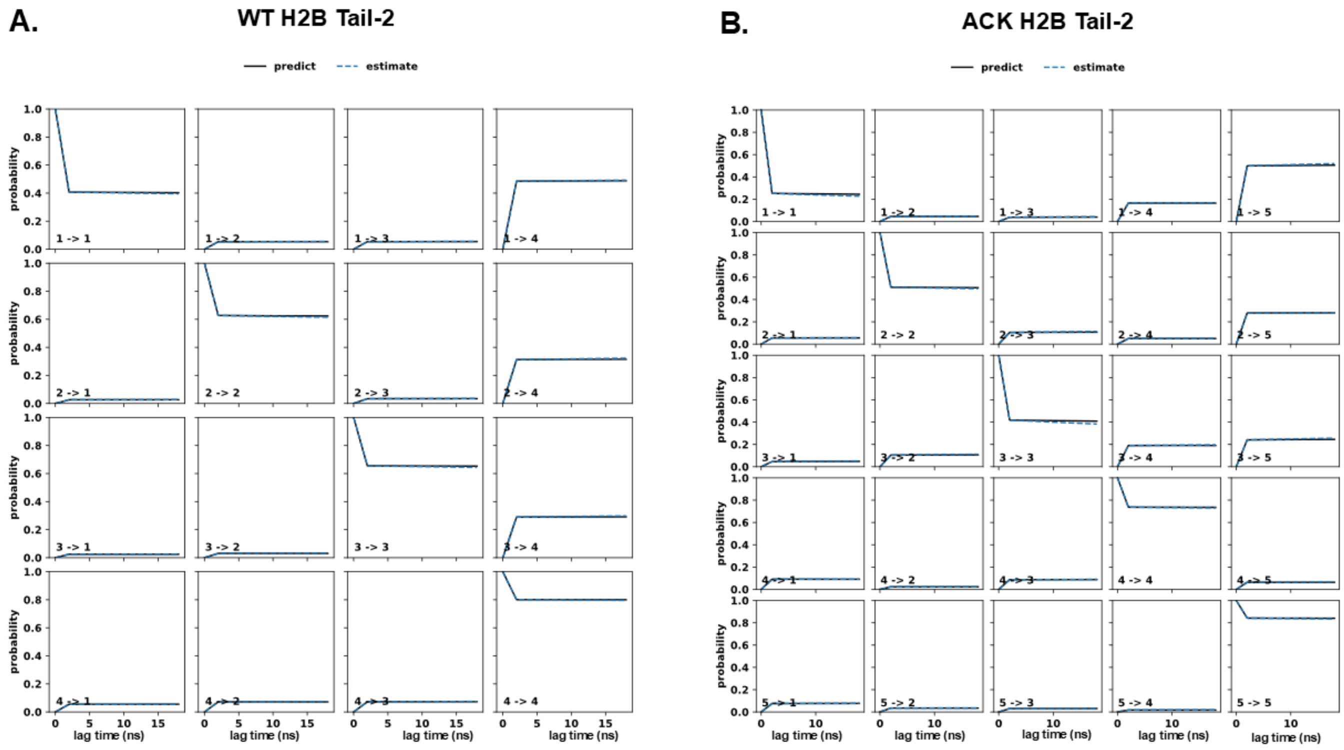


**Fig. S8. Chapman-Kolmogorov (CK) test for MSM validation of ACK H2B Tails-2. (A)** H2B tail-2 and **(B)** ACK H2B tail-2 show four and five metastable states respectively. The predictions from our MSM (blue-dash line) agree well with MSM estimated (solid line) for all states.

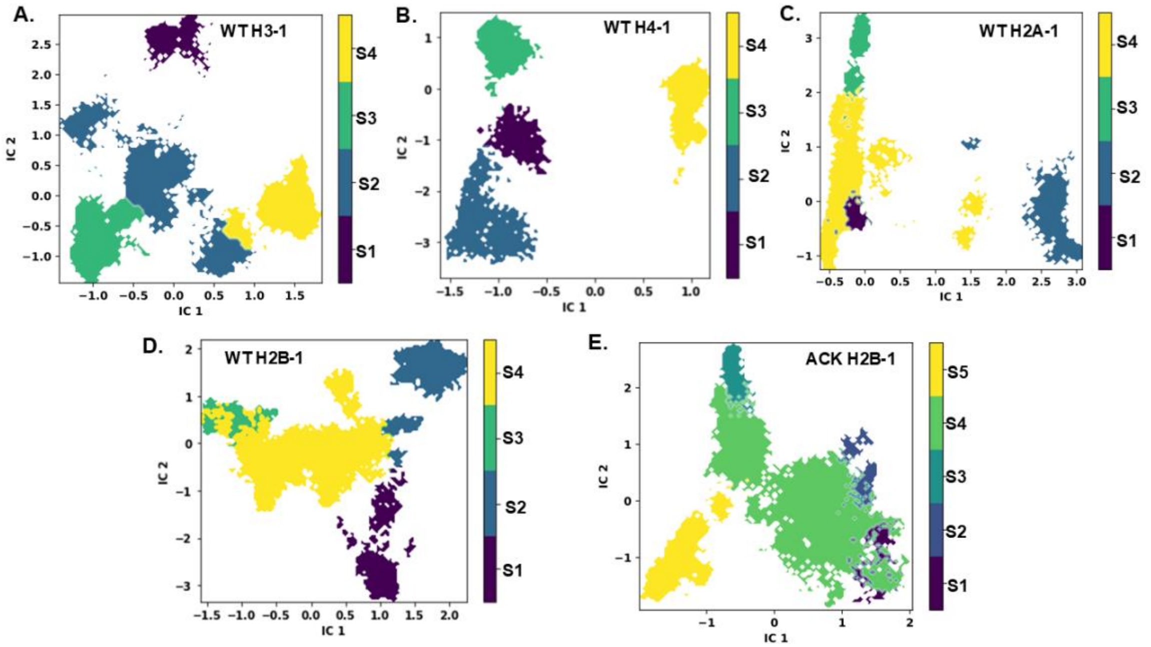


**Fig. S9. Macrostate PCCA+ Clustering Histone Tails-1.** The macrostate clustering visualization projected onto the leading TICA coordinates for visualization of different macrostates for WT **(A)** H3 **(B)** H4 **(C)** H2A **(D)** H2B and **(E)** ACK H2B tails. The metastable macrostates are computed by PCCA+ clustering algorithm within the same TICA projections.

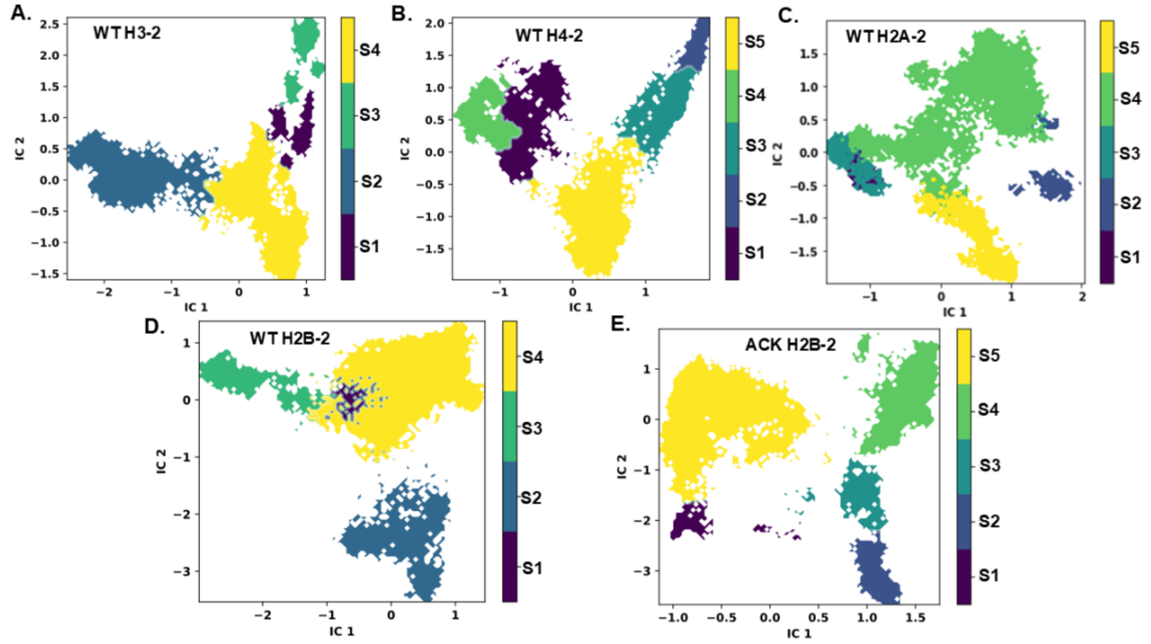


**Fig. S10. Macrostate PCCA+ Clustering Histone Tails-2.** The macrostate clustering visualization projected onto the leading TICA coordinates for visualization of different macrostates for WT **(A)** H3 **(B)** H4 **(C)** H2A **(D)** H2B and **(E)** ACK H2B tails. The metastable macrostates are computed by PCCA+ clustering algorithm within the same TICA projections.

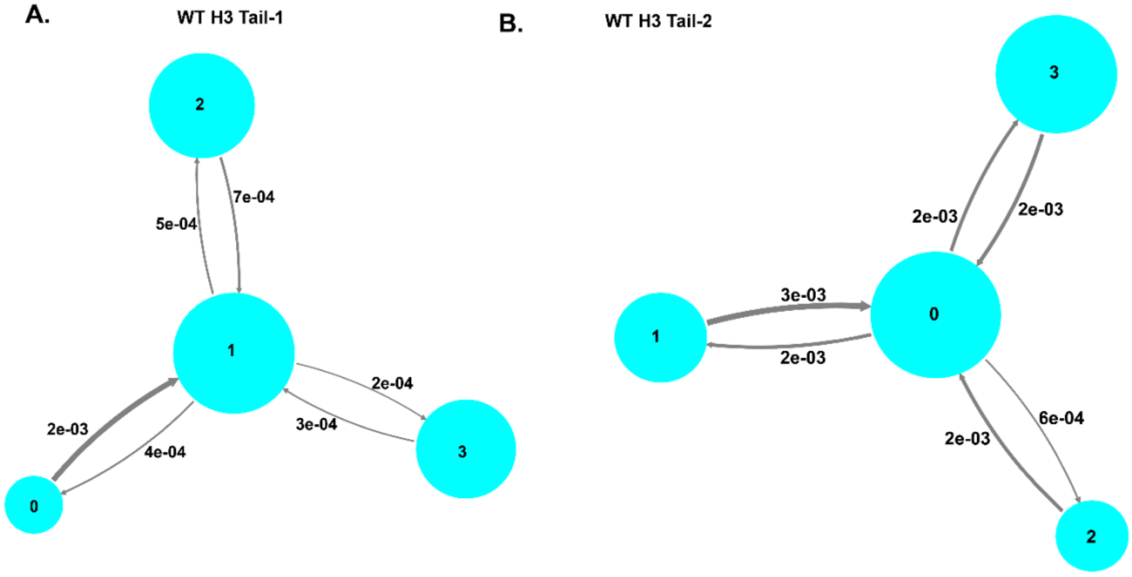


**Fig. S11. MSM Transition probability network plot of H3 Tails. (A)** and **(B)** The MSM network plot connects four macrostates for WT H3 tail-1 and tail-2 represented as cyan circles. The sizes of the circles are proportional to their stationary population for each states. The states are connected through arrows with their transition probability from one state to another is shown next to arrows. The thickness of the arrows is proportional to the amount of flux between the transition of macrostates.

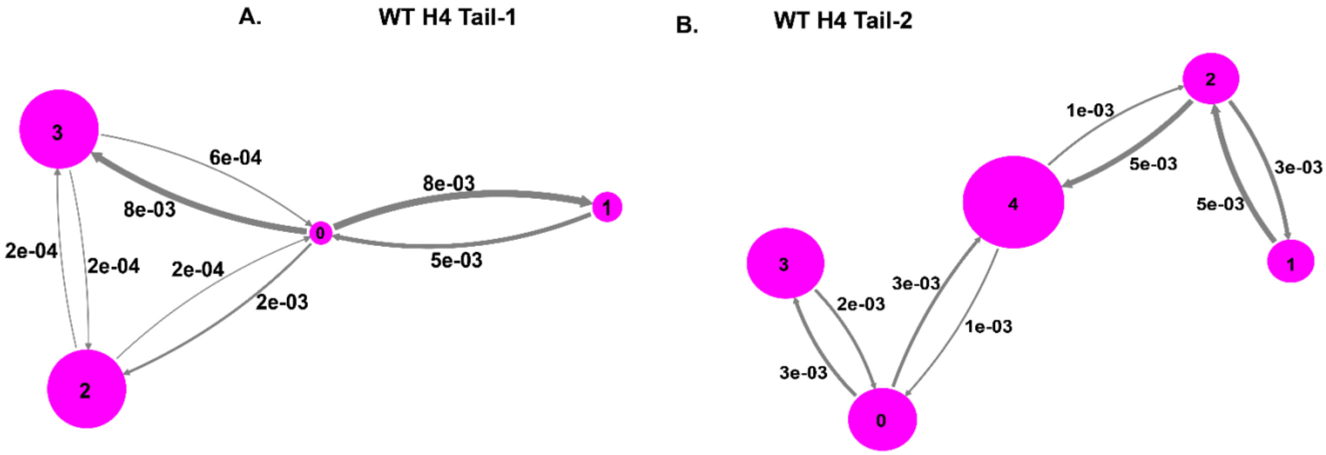


**Fig. S12. MSM Transition probability network plot of H4 Tails. (A)** and **(B)** The MSM network plot connects four and five macrostates for WT H4 tail-1 and tail-2 represented as magenta circles. The sizes of the circles are proportional to their stationary population for each states. The states are connected through arrows with their transition probability from one state to another is shown next to arrows. The thickness of the arrows is proportional to the amount of flux between the transition of macrostates.

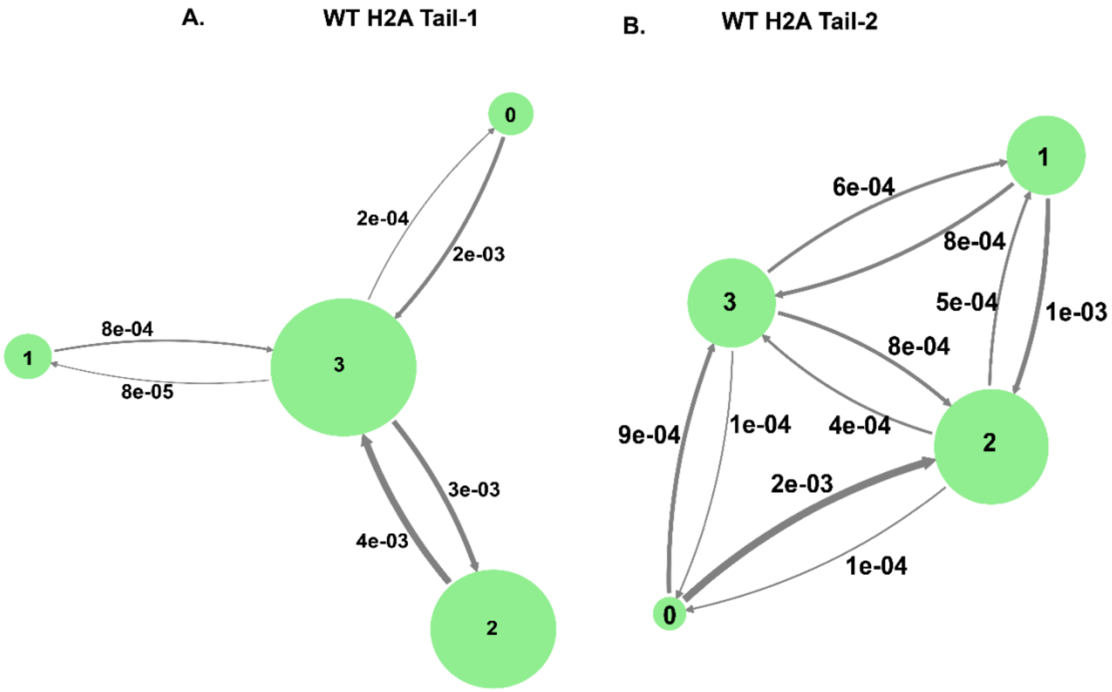


**Fig. S13. MSM Transition probability network plot of H2A Tails. (A)** and **(B)** The MSM network plot connects four macrostates for WT H2A tail-1 and tail-2 represented as green circles. The sizes of the circles are proportional to their stationary population for each states. The states are connected through arrows with their transition probability from one state to another is shown next to arrows. The thickness of the arrows is proportional to the amount of flux between the transition of macrostates.

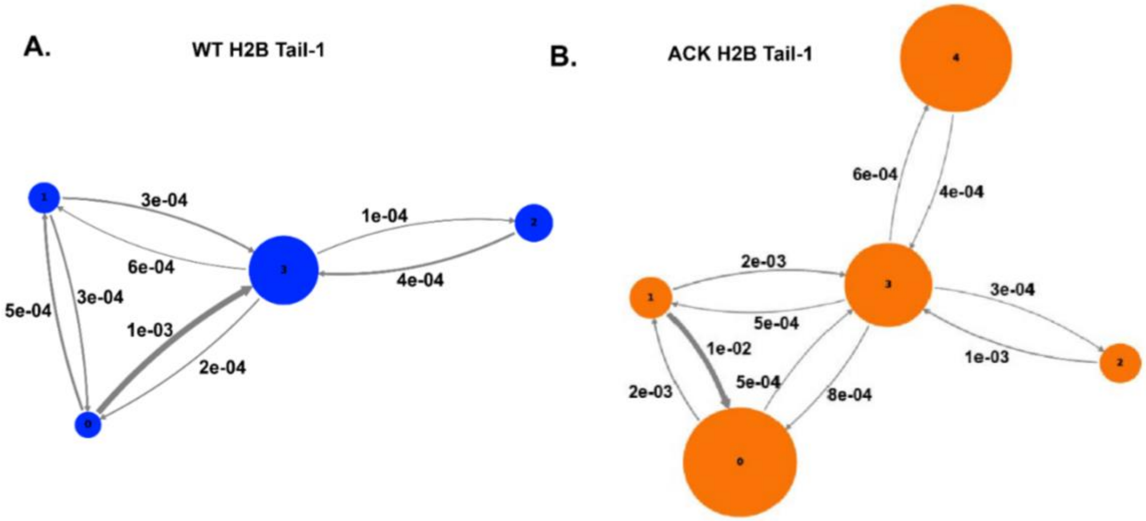


**Fig. S14. MSM Transition probability network plot of H2B Tails. (A)** and **(B)** The MSM network plot connects four macrostates for WT and five for ACK system represented as blue and orange circles respectively. The sizes of the circles are proportional to their stationary population for each states. The states are connected through arrows with their transition probability from one state to another is shown next to arrows. The thickness of the arrows is proportional to the amount of flux between the transition of macrostates.

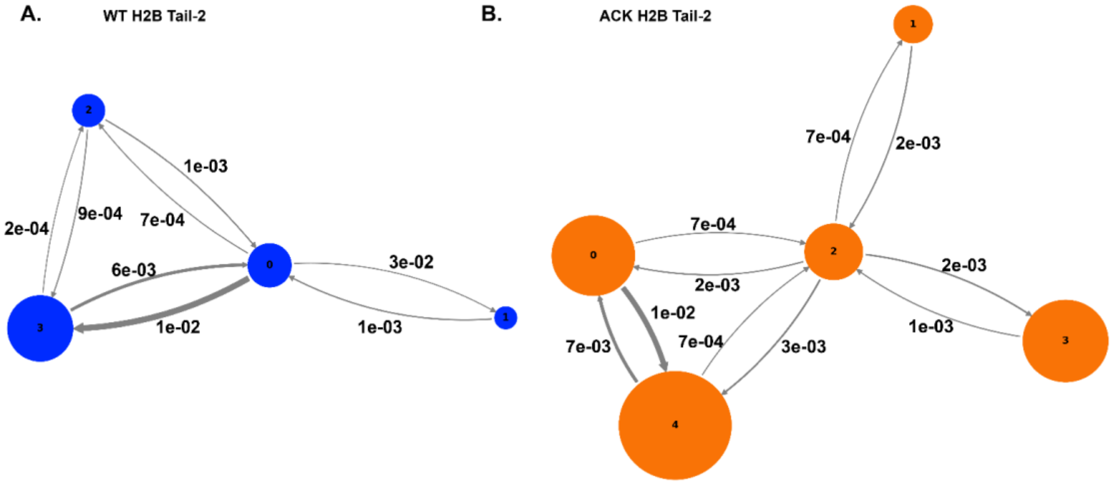


**Fig. S15. MSM Transition probability network plot of H2B Tails. (A)** and **(B)** The MSM network plot connects four macrostates for WT and five for ACK system represented as blue and orange circles respectively. The sizes of the circles are proportional to their stationary population for each states. The states are connected through arrows with their transition probability from one state to another is shown next to arrows. The thickness of the arrows is proportional to the amount of flux between the transition of macrostates.

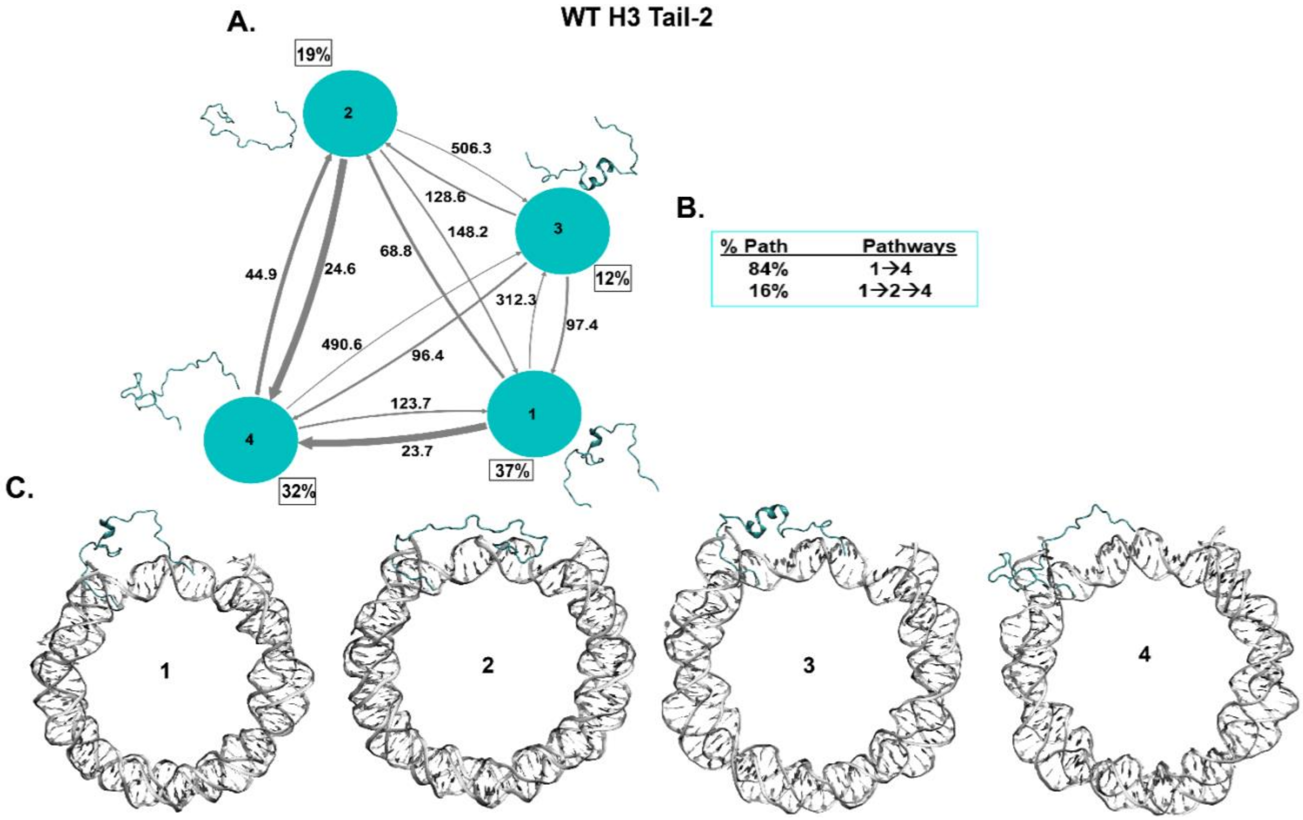


**Fig. S16. The Kinetic network of conformational states of H3 Tail-2. (A)** WT H3 Tail-2 (cyan circles) network plots connect four macrostates of H3 tail-2. The corresponding conformations of each states are shown next to each state for H3 tail-2. The population percentages of each conformations are shown next to each state. The macrostates are connected with arrows. The thickness of the arrows is proportional to the rate of the transition and is labelled with its respective MFPT values in nanoseconds. **(B)** The net flux of the network is obtained from Transition Path Theory (TPT) analysis for H3 tail-2. These TPT calculations show major pathways with their path percentages for H3 tail-2. **(C)** The conformational states of H3 tail-2 with NCP DNA is shown. It shows the position of tail (cyan) at each macrostates with respect to DNA (silver).

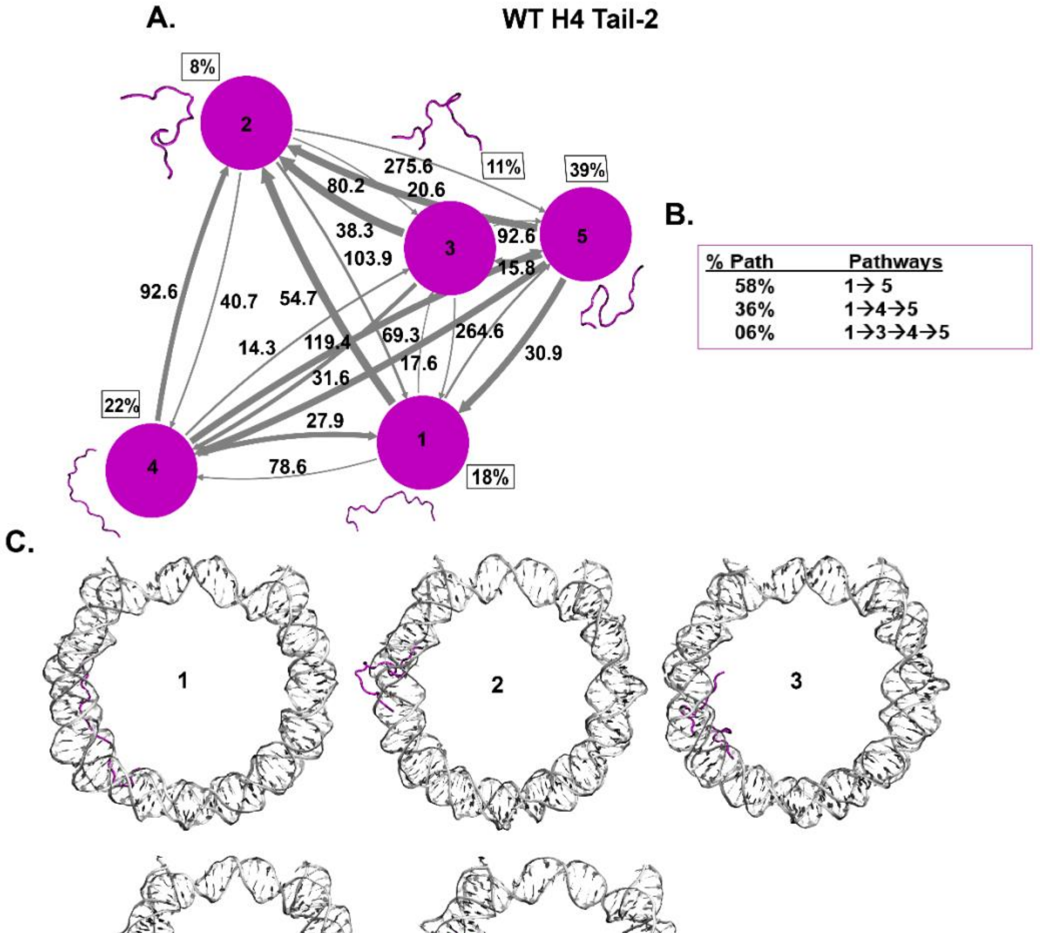


**Fig. S17. The Kinetic network of conformational states of H4 Tail-2. (A)** WT H4 Tail-2 (magenta circles) network plots connect five macrostates of H4 tail-2. The corresponding conformations of each states are shown next to each state for H4 tail-2. The population percentages of each conformations are shown next to each state. The macrostates are connected with arrows. The thickness of the arrows is proportional to the rate of the transition and is labelled with its respective MFPT values in nanoseconds. **(B)** The net flux of the network is obtained from Transition Path Theory (TPT) analysis for H4 tail-2. These TPT calculations show major pathways with their path percentages for H4 tail-2. **(C)** The conformational states of H4 tail-2 with NCP DNA is shown. It shows the position of tail (magenta) at each macrostates with respect to DNA (silver).

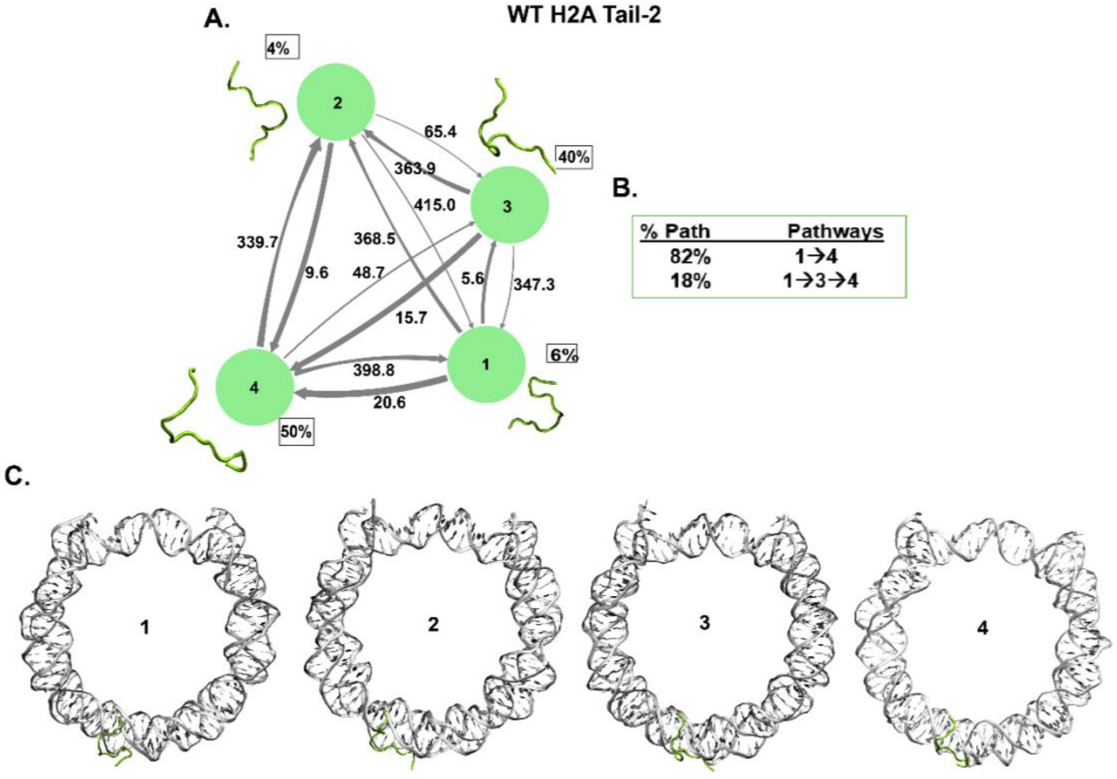


**Fig. S18. The Kinetic network of conformational states of H2A Tail-2. (A)** WT H2A Tail-2 (green circles) network plots connect four macrostates of H2A tail-2. The corresponding conformations of each states are shown next to each state for H2A tail-2. The population percentages of each conformations are shown next to each state. The macrostates are connected with arrows. The thickness of the arrows is proportional to the rate of the transition and is labelled with its respective MFPT values in nanoseconds. **(B)** The net flux of the network is obtained from Transition Path Theory (TPT) analysis for H2A tail-2. These TPT calculations show major pathways with their path percentages for H2A tail-2. **(C)** The conformational states of H2A tail-2 with NCP DNA is shown. It shows the position of tail (green) at each macrostates with respect to DNA (silver).

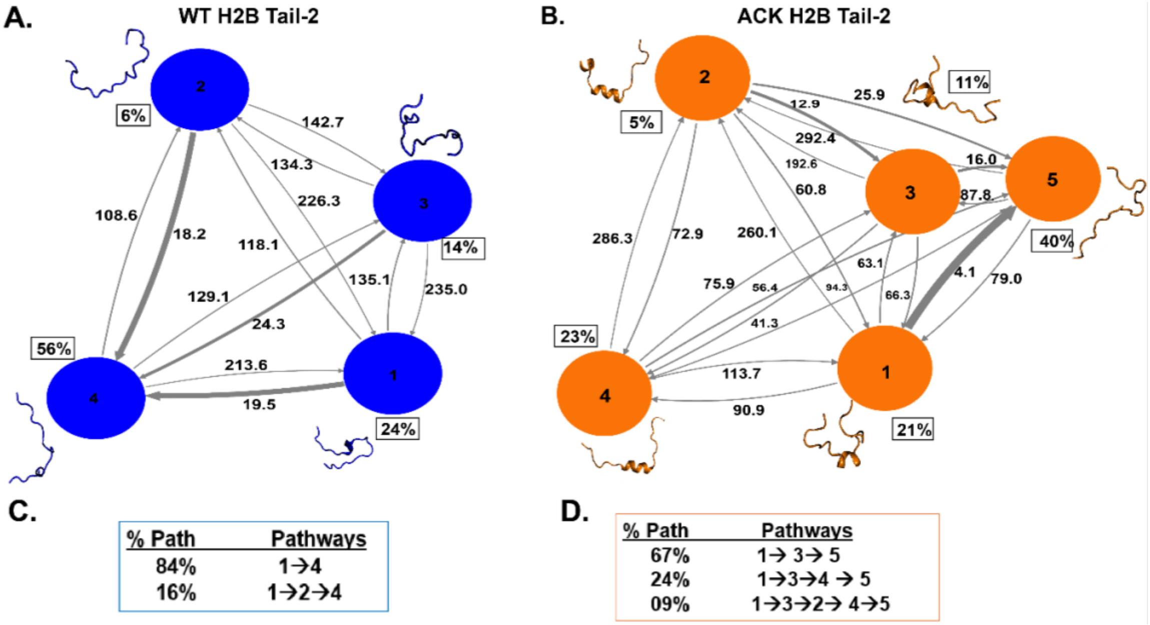


**Fig. S19. The Kinetic network of conformational states of H2B Tail-2. (A)** and **(B)** WT (blue circles) and ACK (orange circles) H2B Tail-1 network plots connect four and five macrostates respectively for H2B tail-2. The corresponding conformations of each states are shown next to each state for both WT (blue) and ACK (orange) H2B tail-2. The population percentages of each conformations are shown next to each state. The macrostates are connected with arrows. The thickness of the arrows is proportional to the rate of the transition and is labelled with its respective MFPT values in nanoseconds. **(C)** and **(D)** The net flux of the network is obtained from Transition Path Theory (TPT) analysis for both WT and ACK H2B tail-2. These TPT calculations show major pathways with their path percentages for both WT and ACK H2B tail-2 systems.

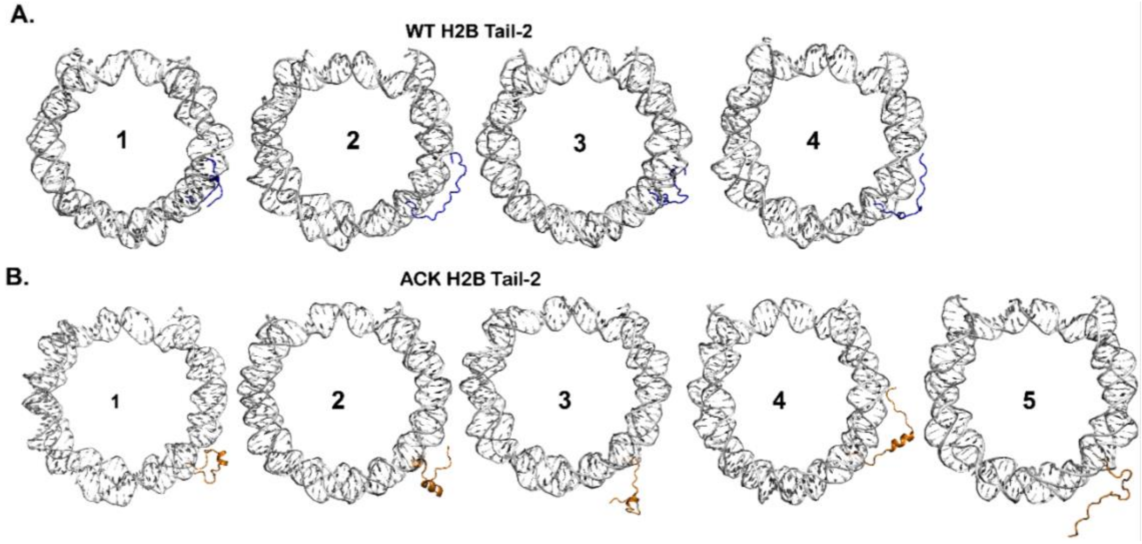


**Fig. S20. The conformational states of H2B Tail-2 with DNA of NCP. (A)** and **(B)** WT (blue) and ACK (orange) H2B Tail-2 conformational states that are shown in Figure 6, the same conformational states of the tail with NCP DNA is shown here. It shows the position of each tail conformations at each macrostates shown earlier with regards to the DNA whether the tail collapsed to the DNA or elongated outwards from the DNA.

**Tables.**

**Table S1.** Summary of NCP systems MD simulation set up box sizes and atoms/ions.

| NCP systems | WT_0.15M (unacetylated) | ACK_0.15M (acetylated) |
|---|---|---|
| **Box Size (Å$^3$)** | 159 x 191 x 112 | 159 x 191 x 112 |
| **No. of atoms** | 444888 | 486492 |
| **No. of Water molecules** | 104740 | 115117 |
| **No. of $Na^+$ ions** | 472 | 508 |
| **No. of $Cl^-$ ions** | 356 | 384 |
| **No. of $Mg^{2+}$ ions** | 14 | 14 |
| **NaCl salt Concentration (M)** | 0.15 | 0.15 |
